# A host-directed virulence factor of *Clostridium perfringens* is modulated by gut commensal strains

**DOI:** 10.64898/2026.03.28.714987

**Authors:** Julia Schumacher, Paolo Stincone, Johanna Rapp, Timo-Niklas Lucas, Carlos Llaca-Bautista, Francesca Barletta, Mirita Franz-Wachtel, Boris Maček, Daniel H. Huson, Lisa Maier, Hannes Link, Daniel Petras, Bastian Molitor

## Abstract

In a healthy host, the residential microbes help regulate the growth of pathobionts, which are common members of the human gut microbiome, preventing them from causing diseases, including infections, under certain conditions. In cases of dysbiosis, this protection may be compromised. Targeted microbiome modulation offers a promising approach to restore healthy conditions in a disrupted community and consequently prevent infections using the natural colonization resistance of the microbiome. Elucidating the interaction mechanisms between microbial species within a microbiome is crucial for understanding how a microbiome can be modulated precisely and effectively to benefit the host’s well-being. Here, we investigated the interactions between the pathobiont *C. perfringens* and human gut commensals on physiological and molecular levels. We found that commensal strains affect *C. perfringens* growth by competing for substrates such as amino acids or a carbon source other than glucose. We further observed that *Bacteroidaceae* strains altered the levels of *C. perfringens* proteins, among others, the host-directed θ-toxin. Our findings reinforce the notion that modulating the composition of the gut microbiome is an effective strategy to prevent infections.

## Background

The pathobiont *C. perfringens* can induce severe diseases in its host^1-3^. Its virulence is linked to its very rapid growth, with a doubling time of below 10 minutes under optimal conditions, and the ability to produce over 20 different strain-dependent toxins or enzymes that are harmful to the host^4^. For energy production, C. perfringens can utilize a variety of different sugars to feed into anaerobic glycolysis or ferment certain amino acids^5^. Unlike other *Clostridia species, C. perfringens* is unable to utilize amino acids via Stickland fermentation to generate ATP, because it lacks key genes for this pathway. However, *C. perfringens* relies on the availability of many amino acids from the environment as it is missing many enzymes for amino-acid biosynthesis^5-7^.

Besides being a common cause of food poisoning, *C. perfringens* can also induce other diseases, including clostridial myonecrosis, necrotic enteritis, enterocolitis, and enterotoxemia^8^. *C. perfringens-* induced food poisoning is usually self-regulating within 24 hours, in part due to the gut microbiome’s colonization resistance, which helps prevent sustained pathogen colonization and overgrowth of pathobionts; therefore, treatment is generally unnecessary beyond maintaining adequate hydration^9,10^. In contrast, other *C. perfringens* infections, including myonecrosis and necrotic enteritis, require immediate action in the form of antibiotic treatment as early as possible, combined with surgical debridement to prevent death. Commonly, penicillin G is administered combined with other broad-spectrum antibiotics such as clindamycin or metronidazole^9^. As observed for other pathogens, the occurrence of infections with antimicrobial-resistant *C. perfringens* is increasing, making treatment of infections sometimes challenging^11-15^. Elimination of pathogens by antibiotic treatment enables resistant strains to further increase in abundance as the sensitive strains are eradicated. In addition, those treatments also impose a strong effect on commensal strains, which can induce dysbiosis^16^.

Even though *C. perfringens* can induce several diseases in the host, it is frequently found to be a normal inhabitant of the intestinal microbiota of animals and humans^17-19^. This indicates that its growth and toxin production are somehow controlled in a healthy environment. Different factors directly influence the members of the microbiota. For example, external factors, such as host genetics, age, diet, geographic location, mode of birth of the host, and intake of medication, can affect the microbiota and also the interspecies interactions between the different members of a microbiota^20^. The importance of those microbial interactions in maintaining homeostasis and preventing pathogen outgrowth is widely appreciated but little is known about those interactions on a molecular level^21,22^. In many cases, correlations between infections or pathogen abundance and the abundance of certain (commensal) strains are reported^23-25^. However, often, it remains unclear whether this is merely a correlation or if enhanced pathogenicity is promoted by surrounding strains or lower numbers of inhibitory strains^26,27^. Microbiome modulation represents a promising approach to restoring a healthy microbiota within a host, for example, through the suppression of pathogens. This would circumvent the collateral damage that many pathogen-targeting drugs have on the commensal strains of the microbiota and offer a new solution to fight multi-drug-resistant pathogens^16^. Due to the complexity of the interplay between microbes, host, environment, and other factors, this seemingly simple approach is still in its infancy and requires further research before it can be translated into clinical use. We need to understand microbe-microbe interactions on a molecular level to precisely modulate the microbiome composition without altering its functionality.

The aim of this study was to investigate how commensal strains of the human gut microbiota interact with *C. perfringens*, using a selection of 19 human gut bacteria (17 commensal strains and two pathobionts). Our data indicate that competition for amino acids dictates, at least in part, the interaction of *C. perfringens* with commensal strains. Further, we provide molecular evidence that *C. perfringens* competes for myo-inositol with *Bacteroidaceae* strains, and that co-cultivation with these same strains alters the abundance of the θ-toxin in *C. perfringens*. Our data highlight the importance of gut commensals in regulating not only pathogen growth but also virulence, which could pave the way for the development of future strategies to fight infections.

## Results

### Interactions with commensal strains prevent long-term colonization of C. perfringens

To investigate how *C. perfringens* is affected by gut commensal strains within a community, we employed a previously described model community, Com18^28^ (**Supplementary Figure 1A**). Com18 contains prevalent abundant members of the human gut microbiota and includes strains from four different bacterial phyla and seven families. In *in vitro* batch cultures, *C. perfringens* becomes the dominant species within this community after 24 hours. However, in a mouse model, while *C. perfringens* is highly abundant after two days, its abundance decreases after six days, and after 57 days, it is no longer detected in the microbiota of these mice^29^. We applied a bioreactor model to investigate *C. perfringens* growth under physiological conditions, in axenic culture or within Com18. Our bioreactor enabled stable growth of *C. perfringens* in axenic culture (**Figure 1A**), but within Com18, *C. perfringens* was almost completely washed out after 116.75 hours (**Figure 1B, Supplementary Figure 1B, bottom panel**). Even though *C. perfringens* was the most dominant strain after 3.25 hours in the bioreactor model, its presence in the community did not have a relevant effect on the relative abundances of the Com18 commensal strains (**Supplementary Figure 1B**).

**Figure 1:**
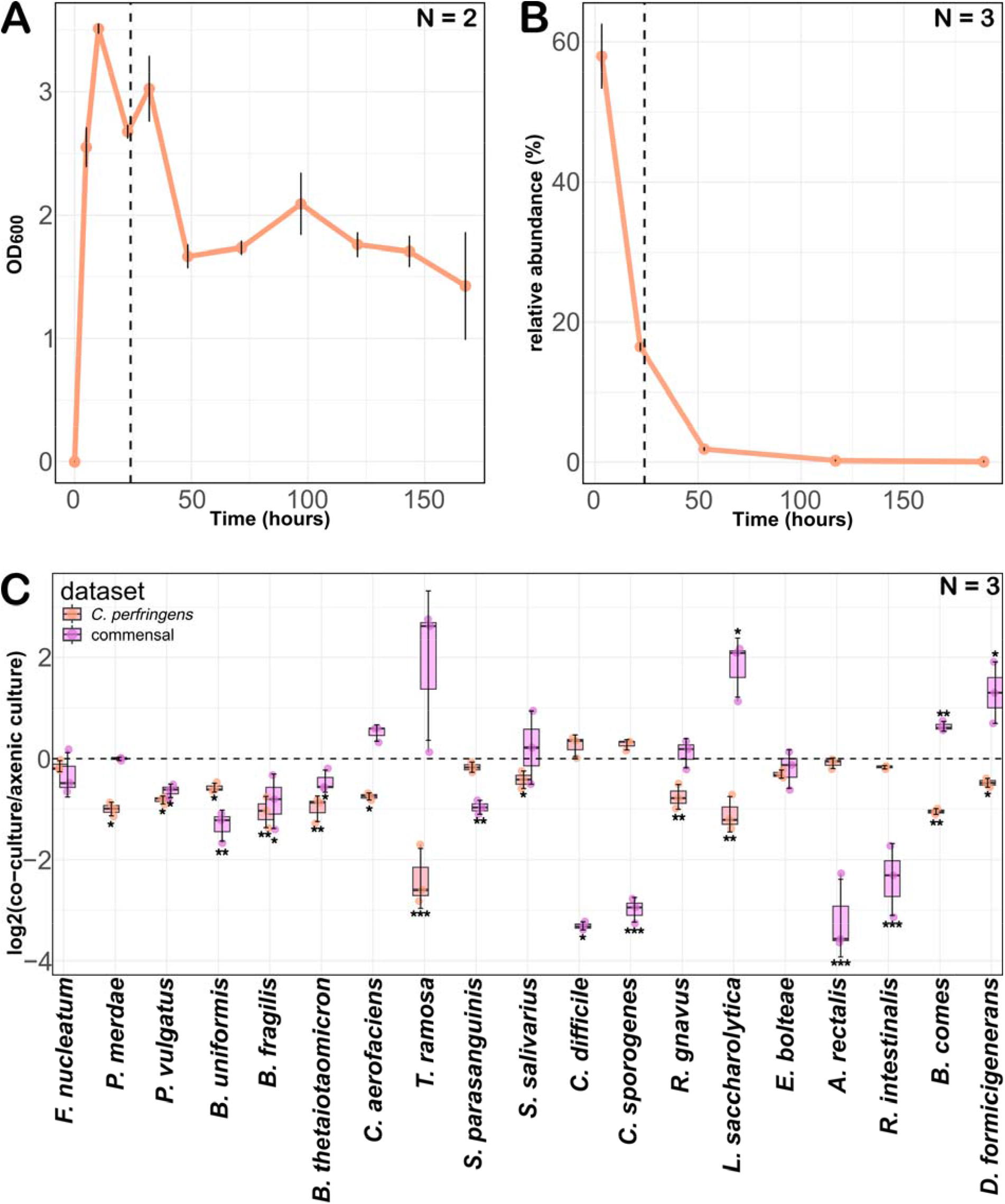
Com18 strains prevent *C. perfringens* long-term colonization. **A:** Growth of *C. perfringens* in monoculture in two independent bioreactors. The first 24 hours were in batch operation before we switched to continuous operation (indicated by the dashed line). Mean value and standard deviation are shown. **B:** Relative abundance of *C. perfringens* in Com18 in three independent bioreactors. After 24 hours (dashed line) the system was switched from batch operation to continuous operation. Mean value and standard deviation are shown. **C:** Log2 fold change of relative abundance of community strains and *C. perfringens* in co-cultures as determined by qPCR. Statistical significance was assessed on relative abundance values using a two-tailed Welch’s t-test. Significance is indicated with *, *p* ≤ 0.05, **, *p* ≤ 0.01, and ***, *p* ≤ 0.001.

To investigate the interaction between each commensal strain and *C. perfringens*, we continued with co-culture experiments in batch. First, we determined the growth of all members of Com18 in axenic culture, and found that *C. perfringens* is the fastest-growing strain to reach the stationary phase, but after 24 hours, all strains reached the stationary phase (**Supplementary Figure 1C**). Thus, we decided to perform follow-up experiments with 24 hours of cultivation. Pairwise co-cultures of *C. perfringens* with each Com18 strain and two further pathobionts, Clostridium *sporogenes* and *Clostridioides* difficile, enabled us to evaluate the abundance of *C. perfringens* and each community strain relative to their abundance in axenic culture (**Figure 1C**).

Based on the relative abundance of *C. perfringens* and each Com18 strain in co-culture, we could define different interaction types (**Supplementary Figure 1D**). An interaction was designated as neutral when the relative abundance of the strains in co-culture was not significantly different from the abundance observed in each axenic culture. In amensal pairwise interactions, one of the strains exhibited a significant inhibitory effect on the other strain, while its abundance remained unaltered. In parasitic interactions, one strain inhibited the other strain to increase in abundance. In contrast, in competitive interactions, the abundance of both strains decreased in co-culture. No mutual interaction was observed, where both co-culture strains would show increased relative abundance (**Figure 1C, Supplementary Figure 1D**). The growth of *C. perfringens* was not sufficiently supported by any of the Com18 strains to result in a higher relative abundance. Conversely, some community strains, namely *Collinsella aerofaciens, Thomasclavelia ramosa, Lacrimispora saccharolytica, Bariatricus comes*, and *Dorea formicigenerans* did exhibit higher relative abundance in co-culture, although this was not significant for *T. ramosa* and *C. aerofaciens* (**Figure 1C**).

### C. perfringens and commensal strains compete for amino acids

To gain a deeper understanding of how bacteria in the co-cultures interact on a molecular level, we analyzed and compared the proteomes of all the co-cultures and single cultures to identify proteins with differential abundance between the co-cultures and the single cultures. We reasoned that these proteins could play a role in the molecular response to the presence of another strain. To reduce the complexity of the data and to get a general overview, we grouped the proteins that were found to be: **(1)** significantly (FC ≥ |2|, adj. P. Val ≤ 0.05) more or less abundant in the co-cultures; **(2)** fully absent in the co-culture; or **(3)** only present in the co-culture, into protein classes according to the categories of the COG database^30^. Within the protein classes that were less abundant or completely absent in the community strains in co-cultures, three main clusters were found to separate the community strains (**Figure 2A**). While one cluster showed only few proteins to be less abundant in the presence of *C. perfringens*, the second and third cluster showed a larger number of less abundant proteins. In the latter two, the largest number of less abundant proteins belonged to the “function unknown” (S) category, highlighting the substantial proportion of microbial proteins in the human gut microbiome with yet uncharacterized functions.

**Figure 2:**
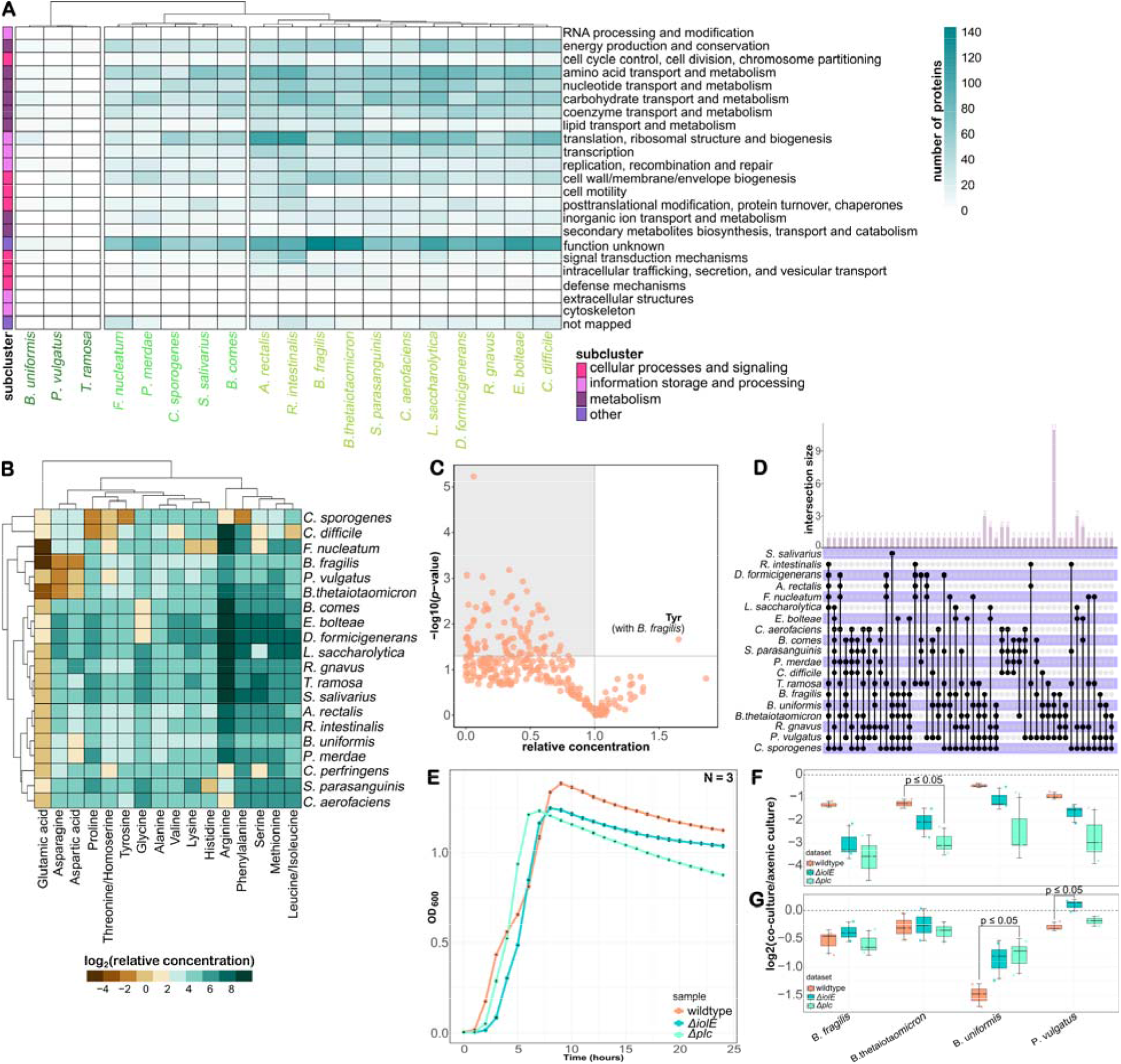
Substrate competition for amino acids and myo-inositol. **A:** Heatmap of community strain proteins that were found to be less abundant in the co-cultures compared to the single culture. Each protein was classified according to the COG database and the number of proteins in each class is visualized in the heatmap. The three main clusters of the heatmap, as given by the Euclidean distance metric, are highlighted by different green font colors of the strain names. Mean values of biological triplicates were used. **B:** Log2 of relative amino-acid concentration normalized to the OD_600_ of each culture (mean value, N = 3). Relative concentrations were calculated by normalizing each metabolite’s intensity to a constant internal standard spiked into all samples. The similarity between strains and metabolites was determined using the Euclidean distance metric. **C:** Measured amino-acid concentration in the co-cultures is plotted relative to the calculated expected amino-acid concentration in the co-cultures. Significantly higher or lower intensities are above the threshold of p < 0.05. (mean value, N = 3). **D:** UpSet plot of *C. perfringens* proteins that were absent or less abundant in the co-culture compared to the single culture. Only proteins that were found in at least four cultures are included. **E:** Growth curve of the two *C. perfringens* deletion strains and the *C. perfringens* wildtype over 24 hours. The OD_600_ was measured every hour. Error bars represent the standard deviation (N = 3). **F:** Log2 of relative growth of *C. perfringens* strains in co-cultures with *Bacteroidaceae* and **G:** Log2 of relative growth of *Bacteroidaceae* strains in co-cultures shown in **F** (N = 3). Statistical significance was assessed using a two-tailed Welch’s t-test.

The clusters identified based on the less abundant proteins of Com18 strains in the presence of *C. perfringens* did not reflect phylogenetic similarities between the strains (**Figure 2A, Supplementary Figure 1A**). Similarly, also the proteins found to be more abundant or only present in the co-cultures with *C. perfringens*, compared to the axenic cultures of the community strains, did not reflect phylogenetic similarities and did not exhibit a discernible clustering of different strains. These data suggest a unique response of each strain to the presence of *C. perfringens* (**Supplementary Figure 2A**). The majority of community proteins found to be less abundant in the presence of *C. perfringens* were found in the “metabolism” subcluster. Thus, we hypothesized that one mechanism of interaction between the Com18 strains and *C. perfringens* is substrate competition. It is known that *C. perfringens* relies on external availability of many amino acids for growth (**Supplementary Figure 2B**). Thus, we asked whether *C. perfringens* could compete for amino acids with Com18 strains. To answer this, we measured the cell-associated amino acids in all single cultures and co-cultures. In total, 16 amino acids were detected in our samples. Glutamine, and tryptophan were not detected, either being absent or below the detection limit of our method. We compared the relative concentration of each amino acid between the axenic cultures of each strain (**Figure 2B**). *C. perfringens* showed a comparatively low concentration of serine, arginine, and threonine/homoserine (**Figure 2B**). These amino acids may be quickly consumed in *C. perfringens* for energy conservation. *C. perfringens* encodes genes for serine and threonine fermentation, as well as for the arginine deaminase pathway, which all result in energy conservation^5,31^. Next, we calculated the expected amino-acid concentration in each co-culture, under the assumption that the two strains in the co-cultures would not affect each other, and compared the expected concentration with the measured concentration of the co-cultures, similar to Medlock, *et al*. (2018): 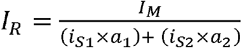 (where *I*_*R*_ = relative metabolite concentration; *I*_*M*_ = measured metabolite concentration in co-culture; *i*_*S1*_ = measured metabolite concentration of community strain in single culture; *a*_*1*_ = abundance of community strain in co-culture; *i*_*S2*_ = measured metabolite concentration of *C. perfringens* in single culture; *a*_*2*_ = abundance of *C. perfringens* in co-culture) (**Figure 2C**). The concentration of most amino acids in the co-cultures was lower than expected (**Figure 2C**). This could be the result of amino-acid cross-feeding in the co-cultures, where one strain consumes the amino acids synthesized by the other strain, thus reducing the overall concentration, or it could be the consequence of a regulatory process.

In the co-culture of *C. perfringens* with B. fragilis, tyrosine showed a significantly higher intensity than expected (**Figure 2C**). Investigating the differentially abundant proteins in the co-culture as compared to their single cultures revealed a decreased abundance of a B. fragilis putative aspartate aminotransferase (Q5LEZ7) (**Supplementary Figure 2C**). This enzyme is predicted to bi-directionally convert 4-hydroxypyruvate to tyrosine. In the presence of *C. perfringens, B. fragilis* possibly reduces the conversion towards 4-hydroxypyruvate.

By investigating the differentially abundant proteins of *C. perfringens* in the co-cultures, we found that the proteomic response of *C. perfringens* did not correlate with the phylogenetic relatedness of the community strains, as even closely related strains elicited distinct responses (**Supplementary Figure 3A-B**). However, some *C. perfringens* proteins were differentially abundant in the presence of all tested *Bacteroidaceae* strains (**Figure 2D, Supplementary Figure 3C, Supplementary Figure 4A-D**). Among them, DapH (UniProt: Q0TP51), a protein involved in lysine biosynthesis, showed increased abundance in the co-cultures with all four *Bacteroidaceae*, although this increase was not specific to these conditions. By contrast, 11 *C. perfringens* proteins were exclusively less abundant in all four *Bacteroidaceae* co-cultures (**Figure 2D, Supplementary Figure 3C**).

**Figure 3:**
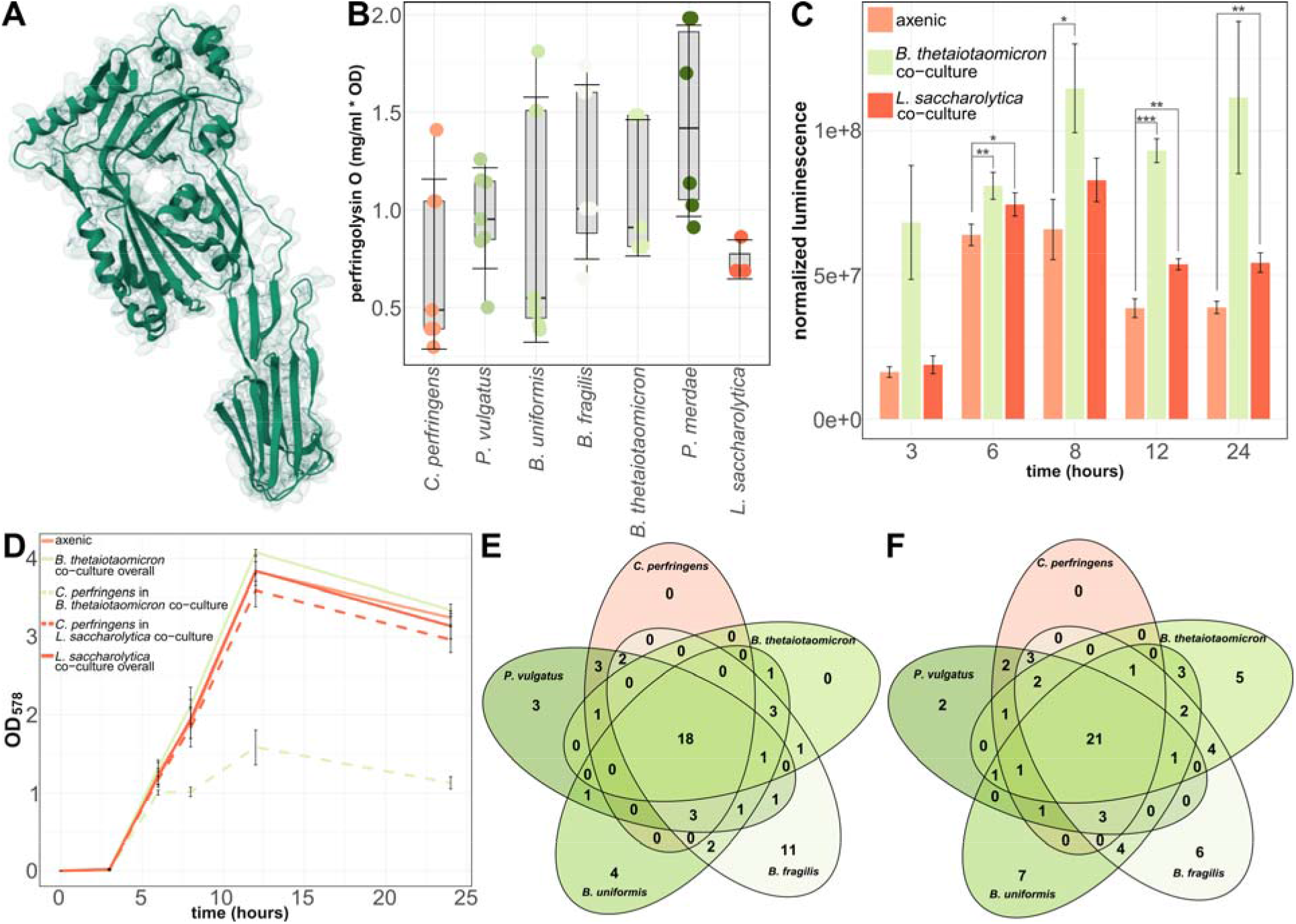
Bacteroidaceae strains modulate *C. perfringens* virulence. **A:** Ribbon model of PFO structure determined by X-ray crystallography^40^ (PDB, 1M3I) from RCSB PDB^41^, deposited by Rossjohn, *et al*. (2004). **B:** Concentration of PFO in mg/mL per OD_600_ of *C. perfringens* (determined by qPCR). Technical replicates of biological triplicates are shown. For the *P. merdae* co-culture only two biological replicates were analyzed and only one biological replicate for the *L. saccharolytica* co-culture. **C:** P_*pfo*_ promoter activity per OD_600_ of *C. perfringens* (determined by qPCR) in single culture (axenic), or in co-culture. The *p*-values were determined through a two-tailed Welch’s t-test. Significance is indicated with *, *p* ≤ 0.05, and **, *p* ≤ 0.01.

### C. perfringens competes for myo-inositol with Bacteroidaceae

Within these conserved *C. perfringens* proteins, we identified IolE (UniProt: Q0TUZ0), an enzyme involved in myo-inositol catabolism (**Supplementary Figure 4A-D**). Two further *C. perfringens* proteins involved in myo-inositol catabolism were less abundant in the presence of the *Bacteroidaceae*. Those were IolB (UniProt: A0A0H2YNH4) and IolG_2 (UniProt: A0A0H2YR21). Despite the absence of significant differences in their abundance across all four Bacteroidaceae co-cultures, the distinct regulatory responses observed only in the co-culture conditions motivated us to further examine these proteins. (**Supplementary Figure 4A-D**). It has been demonstrated that numerous *Bacteroides* species are capable of synthesizing inositol lipids, which possess a myo-inositol phosphate headgroup^33^. Moreover, it has been shown that *C. perfringens* can utilize myo-inositol as an alternative carbon source in the absence of glucose^34^. Although we did not supplement myo-inositol in our mGAM medium, the medium includes yeast extract, which potentially contains myo-inositol. Thus, we hypothesized that *C. perfringens* and the *Bacteroidaceae* strains compete for myo-inositol in the co-cultures, and that *C. perfringens* showed a decrease in the abundance of proteins required for myo-inositol catabolism because their expression depends on the availability of myo-inositol^34^. Indeed, if *Bacteroidaceae* strains use environmental myo-inositol for their lipid metabolism, the concentration of myo-inositol is expected to decrease in co-culture as compared to *C. perfringens* single culture. We anticipated that in case of competition for myo-inositol between *Bacteroidaceae* strains and *C. perfringens*, a *C. perfringens* deletion strain unable to catabolize myo-inositol as alternative carbon source would have a disadvantage compared to the wildtype *C. perfringens* in single culture and in co-culture. Alternatively, *C. perfringens* could acquire myo-inositol through the activity of its α-toxin, which is the phospholipase Plc, and which has an astonishingly broad substrate range^35^. Thus, the α-toxin could enable *C. perfringens* to hydrolyze lipids, making their head groups, including myo-inositol, available for energy metabolism. In this scenario, a *C. perfringens Δplc* would have a fitness disadvantage compared to wildtype *C. perfringens*. To test these two assumptions, we constructed a *C. perfringens* Δplc and a ΔiolE deletion strain. To validate the CRISPR-Cas9 mediated *Δplc* deletion, we developed GeneGone, a lightweight web application for general validation of gene deletions from sequencing data. The tool aligns reads from mutant strains to a wild-type reference genome and computes coverage profiles across the targeted locus. A deletion is indicated by a pronounced drop in coverage over the target region compared to the average coverage in flanking regions. Thresholds are automatically adjusted depending on sequencing read type (Illumina vs. long-read), ensuring reliable detection across different datasets. Applying GeneGone to our *C. perfringens Δplc* mutant confirmed clean deletion of the targeted gene.

Next, we investigated the growth behavior of the *C. perfringens* Δplc and the *C. perfringens ΔiolE* deletion strain in single culture (**Figure 2E**) and in co-cultures with the four *Bacteroidaceae* strains (**Figure 2F**). Both deletion strains showed a similar growth rate as the *C. perfringens* wildtype but reached the stationary phase at a lower OD_600_, which was also reflected in a lower OD_600_ after 24 hours (**Figure 2E**). The wildtype showed a biphasic growth behavior that was not observed in the deletion strains, which could indicate that *C. perfringens* switched to another substrate during exponential growth. However, this biphasic growth was not seen in all *C. perfringens* growth experiments (**Supplementary Figure 1C**). In the co-cultures, the fitness of the community strains (compared to the single culture) did not change significantly between the wildtype *C. perfringens* and the deletion strains, except for *P. vulgatus* and *B. uniformis*, which showed increased fitness in the presence of the *C. perfringens ΔiolE* or the *C. perfringens Δplc* strain (**Figure 2G**). Compared to the wildtype, both *C. perfringens* deletion strains exhibited a tendency toward reduced fitness in the presence of the *Bacteroidaceae* relative to their single culture. This supported our hypothesis that *C. perfringens* competes for myo-inositol in those co-cultures and that IolE is required for myo-inositol catabolism. Since the *C. perfringens Δplc* mutant showed a more pronounced fitness loss than *ΔiolE* in co-cultures, Plc is likely to be relevant for additional functions, not necessarily related to myo-inositol metabolism.

### Bacteroidaceae strains modulate C. perfringens virulence

Among the less abundant proteins of *C. perfringens* exclusive to co-cultures with *Bacteroidaceae*, we also identified the θ-toxin perfringolysin O (PFO) (**Figure 2D, Figure 3A, Supplementary Figure 4A-D**). PFO plays a substantial role during *C. perfringens* infection by inducing the formation of pores in the membrane of host cells, which ultimately results in cell lysis. It has been observed that the toxin specifically targets the cholesterol moieties in the membrane, and as a result, it is believed to not affect prokaryotic cells^36^.

Since our proteome analysis only reflected the intracellular proteins, we analyzed the PFO concentration in the supernatant of the *C. perfringens* axenic culture and of co-cultures with the four *Bacteroidaceae*, as well as *P. merdae* and *L. saccharolytica* co-cultures as control (**Figure 3B**). We found slightly elevated concentrations of PFO normalized to OD_600_ of *C. perfringens* in the supernatant when a *Bacteroidaceae* strain or *P. merdae* was present. The intracellular toxin concentration was not comparable with the concentration in the supernatant. Possibly, PFO secretion is induced in the presence of the *Bacteroidaceae* but the production remains unchanged. This would decrease the intracellular PFO concentration. Decreased levels of PFO could also be explained by a decreased toxin-gene expression. To test these two possibilities, we studied the P_*pfo*_ promoter activity with a luciferase reporter system in axenic culture and in co-culture with *B. thetaiotaomicron* or *L. saccharolytica* as a control, at different growth phases (**Figure 3C**). The promoter activity in the axenic culture increased during the exponential phase, decreased again in the late exponential phase (after 12 hours), and remained lower in the stationary phase (24 hours). Instead, in the co-cultures with *B. thetaiotaomicron* and *L. saccharolytica*, the promoter activity was elevated at all time points. Interestingly, this inversely reflected the OD_600_ of *C. perfringens* in these cultures (**Figure 3D**). The lower the *C. perfringens* OD_600_, the higher the P_*pfo*_ promoter activity at each time point. These findings seemed to contradict the results of the proteome analysis but could be explained by increased *pfo* expression coupled with increased PFO secretion or degradation.

To investigate the proteomic changes in *C. perfringens* exclusive to co-cultures with *Bacteroidaceae*, we asked whether the *Bacteroidaceae* strains had a similar metabolome that could potentially correlate with the conserved *C. perfringens* proteome response. Thus, we measured the cell-associated metabolomes and the exometabolome of *P. vulgatus, B. uniformis, B. fragilis, B. thetaiotaomicron*, and *C. perfringens* from the same axenic cultures used to analyze the proteome using untargeted liquid chromatography-tandem mass spectrometry (LC-MS/MS). We found that the main metabolite class, as predicted by CANOPUS, in all five cultures, cell-associated and in the supernatant, was carboxylic acids and derivatives (**Supplementary Figure 5, Supplementary Figure 6**)^37^. In the cell-associated metabolomes, this metabolite class accounted for roughly 80% to 85% of all measured metabolites (**Supplementary Figure 5**). In the exometabolomes, it accounted for < 50% of all metabolites (**Supplementary Figure 6**). After the carboxylic acids and derivatives, the most abundant metabolites associated with *C. perfringens* were organonitrogen compounds, organooxygen compounds, and fatty acyls (**Supplementary Figure 5**).

At first glance, the cell-associated- and exometabolomes of all five strains looked very similar. Therefore, we looked for metabolite classes found in all *Bacteroidaceae* strains but not in *C. perfringens* (**Figure 3E-F**). While most metabolite classes were found in all five strains, we found one metabolite class in the cell-associated metabolomes (sphingolipids) (**Figure 3E**), and one metabolite class in the exometabolome (quinolones and derivatives) (**Figure 3F**) unique to the *Bacteroidaceae* strains but not present in *C. perfringens*. Sphingolipids are commonly found in eukaryotic cell membranes, but some bacteria, including *Bacteroidaceae*, can include them in their membranes^38^. While some bacteria were found to produce quinolone derivatives, this was not reported for Bacteroidaceae^39^. Even though this has not been investigated before, it is unlikely that our medium contained quinolones. Additionally, in a metabolite profiling approach via flow-injection MS (FI-MS) screen of cell-associated metabolomes of all axenic- and co-cultures, m/z-features annotated as different sphingolipids were detected (**Supplementary Figure 7**). Most sphingolipids here showed the highest intensity in the *Bacteroidaceae* single cultures. Except for N-acetylsphingosine, which showed a low intensity in *C. perfringens*, no other sphingolipids were detected in the *C. perfringens* metabolome. In most co-cultures, the intensity was lower than in the single cultures, which could also be due to lower strain abundance in the co-cultures. Overall, our data demonstrated altered PFO levels in the presence of commensal *Bacteroidaceae* strains. While the molecular pathways responsible for this modulation remain to be elucidated, we observed sphingolipids and quinolones to be more prevalent in *Bacteroidaceae* strains compared to other Com18 strains and *C. perfringens*. The role of these compounds in regulating *C. perfringens* virulence has not been addressed and warrants further investigation.

The error bars present the standard deviation (N = 3). **D:** Growth of ***C. perfringens*** in single culture (axenic), or in co-cultures and overall growth of the co-cultures. The OD_600_ of ***C. perfringens*** in the co-cultures was determined by qPCR. The error bars indicate the standard deviation (N = 3). **E:** Venn diagram of metabolite classes found in the cell-associated metabolomes and **F:** in the exometabolomes.

## Discussion

In this study, we describe the interactions between prevalent and abundant human gut commensal strains and *C. perfringens* on different levels. We examined the interactions on a physiological level, by growing each commensal strain in a co-culture with *C. perfringens* and found that under our conditions, none of the commensal strains enhanced *C. perfringens* growth, but either inhibited *C. perfringens* or did not influence its growth. Accordingly, *C. perfringens* could not stably grow within Com18 in a chemostat. Another study investigating pairwise interactions between bacterial members of a different synthetic gut community also reported a lack of mutualistic interactions^43^. The authors showed that the interactions changed with different media compositions. Similarly, we expect that the interactions between commensal strains and *C. perfringens* will be different in other environments, for example, different media. Our mGAM medium is a complex and rich medium, supporting the growth of a broad range of bacteria. The use of minimal media could reveal explicit metabolic dependencies in co-cultures. We further studied the interspecies interactions on a molecular level through proteome and metabolome analyses. With the metabolome data, we focused first on the amino-acid requirements of *C. perfringens*, which relies on the environmental availability of many amino acids^5-7^. It is known that amino-acid auxotrophic *E. coli* strains can grow in the presence of certain non-auxotrophic strains when the required amino acid is not supplemented in the medium in a process known as cross-feeding^44^. Recent work has shown that the auxotrophic *E. coli* strains are not limited to taking up amino-acids directly, but can also use metabolic intermediates of amino-acid biosynthesis pathways, produced by other strains, to synthesize the required amino acids^45^. *C. perfringens* encodes genes associated with all amino acid biosynthetic pathways. However, many of these pathways appear incomplete, containing only a subset of the required enzymes^5^. It is, therefore, likely that *C. perfringens* can grow when the necessary intermediate metabolites were externally available. We mostly observed lower amino-acid intensities in the co-cultures with *C. perfringens* than expected from the intensities in the single cultures. This could be the consequence of *C. perfringens* consuming not only amino acids but also the intermediates of amino-acid biosynthesis produced by other strains. By depleting these precursors, *C. perfringens* may limit amino acids synthesized in the community, leading to overall lower amino-acid levels in co-cultures. Substrate competition between *C. perfringens* and Com18 strains was not only seen for amino acids but also for myo-inositol. In this case, the competition was exclusively observed in co-cultures of *C. perfringens* and the Bacteroidaceae strains. By targeting a single gene, *iolE*, in *C. perfringens*, we could reduce its fitness by preventing the strain to use myo-inositol. This demonstrates that the use of alternative substrates like myo-inositol, which seems to be of minor importance in axenic cultures, can be crucial for the fitness of a strain in co-cultures.

Our data highlights the importance of *Bacteroidaceae* strains in regulating *C. perfringens* virulence, as PFO levels were changed. However, we did not investigate whether *pfo* expression is co-regulated with the *C. perfringens*-carbon metabolism. Ohtani, *et al*. (2010) found that the expression of the myo-inositol operon is positively regulated by the VirR/VirS system. Despite this, it was shown that *pfo* expression is regulated by two different promoters, and while the main promoter seems to be induced by VirR/VirS, the other promoter is independent of this system^47^. Our luciferase reporter system included both promoters, and therefore, did not allow for differentiation between the two. Further analysis of the *pfo* mRNA levels in *C. perfringens* under different conditions, or two separate reporter systems for the two promoters could reveal which promoter activity is altered in the presence of the *Bacteroidaceae*. The myo-inositol operon was shown to be regulated by myo-inositol and glucose^34^: **(1)** Myo-inositol induces the expression of the operon *via* VirR and a regulatory RNA; **(2)** In the same study, it was stated that glucose repressed the operon if present in an equal or higher concentration than myo-inositol. It was further stated that myo-inositol did not induce toxin production, but no data was presented, and it is not evident which toxins were evaluated^34^. Therefore, the influence of myo-inositol and glucose on PFO production should be further studied, which is now possible with the deletion strains generated in our study.

*Bacteroidaceae* strains are able to synthesize sphingolipids and inositol lipids^33,48^. Both lipids are found in many eukaryotic cell membranes but are comparably rare in prokaryotic cells. Research on *Bacteroidaceae-*derived inositol- and sphingolipids has focused on the relevance of those lipids for host signaling^38,48-50^. Little is known about their role in signaling within microbiota, but our metabolite profiling approach *via* FI-MS suggests that this metabolite class is relevant for *C. perfringens* virulence. The data presented here provide a valuable foundation for further investigations of *C. perfringens* interactions in the human gut, but limitations of the study should also be acknowledged. Both, our proteome and metabolome analyses widely rely on annotations from public databases or *de-novo* annotations from software. A metabolite can only be fully validated by confirming the structure and metabolite assignment using a reference standard, and the function of a detected protein can be validated only experimentally. Additionally, in our experimental design, we selected 24 hours as the end time point to examine the interactions between *C. perfringens* and human gut commensals because the breadth of systems biological samples did not allow us to examine intermediate time points. In future experiments, this would have the potential to give further insight on the growth kinetics of the strains and on the nature of the molecular interactions.

In summary, this study illustrates the importance of substrate competition between *C. perfringens* and commensal strains in regulating the fitness of the pathobiont in co-cultures. We showed that the *Bacteroidaceae* strains not only affect *C. perfringens* growth but also have an impact on its virulence. Even though the signaling cascade regulating toxin production in *C. perfringens* in response to the *Bacteroidaceae* strains remains to be elucidated, our study provides the basis for future strategies to exploit microbiome interactions to control pathogen virulence.

## Methods

### Strains and growth conditions

The strains used in this study are listed in **Supplementary Table 1**. All gut microbiome strains were grown in modified GIFU anaerobic medium (mGAM) (HyServe GmbH & Co.KG, Product Code: 1005433-001) inside a Coy chamber (95 vol-% N_2_ and 5 vol-% H_2_) (Coy Laboratory Products Inc.). Cultures were incubated at 37°C, non-shaking. All strains were inoculated from glycerol stocks into liquid medium and transferred again after 24 hours before they were used for experiments the following day. For cultivation on plates, 1.5 weight-% agar (Carl Roth, Agar-Agar Kobe I, CAS 9002-18-0) was added to the mGAM medium. All media were prepared aerobically but transferred into the Coy chamber at least 24 hours prior to usage for pre-reduction. If necessary, the liquid medium and plates were supplemented with 15 μg/mL thiamphenicol (Sigma-Aldrich, CAS-15318-45-3), 5 μg/mL erythromycin (Thermo Scientific, CAS 114-07-8), 50 μg/mL 5-aminolevulinic acid hydrochloride (Sigma-Aldrich, CAS 5451-09-2), or 5 mM theophylline (Sigma-Aldrich, CAS 58-55-9). Theophylline was dissolved in DMSO (Fisher Scientific, CAS 67-68-5).

For all co-culture assays, all strains were inoculated from cryo stocks into 3 mL mGAM medium and incubated overnight. The next day, all cultures were diluted 1:300 in 3 mL fresh mGAM and incubated again overnight. The following day all single- and co-cultures were set up in 25 mL mGAM medium with a starting OD_600_ of 0.001 per strain (therefore, co-cultures had a starting OD_600_ of 0.002 and single cultures a starting OD_600_ of 0.001). If not indicated otherwise, the cultures were incubated for 24 hours before being further analyzed.

*E. coli* strains used in this study were grown aerobically in Luria Broth (LB, contained per liter: 10 g peptone, 5 g yeast extract, 10 g NaCl) or on LB agar plates (1.5 weight-%) at 37°C. Liquid cultures were incubated shaking at 150 rpm. If required, the medium was supplemented with 30 μg/mL chloramphenicol (Carl Roth, CAS 56-75-7) or erythromycin (250 μg/mL, liquid; 400 μg/mL, plates) (Thermo Scientific, CAS 114-07-8). E. coli ST18 is auxotrophic for 5-aminolevulinic acid, therefore, its medium was supplemented with 50 μg/mL 5-aminolevulinic acid hydrochloride (Sigma-Aldrich, CAS 5451-09-2).

### Bioreactor handling and operating conditions

The bioreactor setup and operating conditions were as described in Schumacher, et al. (2025). In brief, the working volume was 250 mL, the temperature was set to 37 °C (± 0.5 °C), and the pH was maintained at 7 (± 0.05; hysteresis = 0.01). The pH was constantly monitored and adjusted using 0.5 M HCl and NaOH. To maintain anaerobic conditions, the bioreactors were continuously sparged with N_2_ gas at a rate of approximately 2.5 mL min^-1^. Each microbe was transferred and grown twice in single culture before inoculation of the bioreactors. The starting OD_600_ was always 0.01 and all strains contributed equally to the starting OD_600_. After 24 hours in batch operation, the bioreactors were set to continuous operation with a hydraulic retention time (HRT) of 24 hours, resulting in a flow rate of 0.1736 mL per minute. Samples for OD_600_ measurements were taken at least every second day and samples for sequencing or qPCR analysis were taken as indicated in the results.

### 16S rRNA gene amplicon sequencing

The 16S rRNA gene amplicon sequencing for the determination of the strain abundances in our multiple-bioreactor system (MBS) was executed and analyzed as described in a previous study^28^. Genomic DNA from each sample was extracted using the DNeasy UltraClean 96 Microbial Kit (Qiagen 10196-4). Primers 515F and 806R were used for amplification, and the library preparation and sequencing were performed by the NGS 658 Competence Center NCCT (Tübingen, Germany) on an Illumina MiSeq device using the v2 sequencing kit. The computational processing of the obtained sequences was performed as described before^28^.

### Relative abundance analysis of strains by qPCR with species-specific primers

To determine the relative abundance of the strains in co-cultures we designed species-specific primers (**Supplementary Table 2**). Strain-specific genes were determined using the MetaPhlAn 3 database (http://segatalab.cibio.unitn.it/tools/metaphlan3/) and primers were designed manually within those genes^51^. The primers were designed to match the following parameters: amplicon size between 184 bp and 257 bp, primer length between 24 bp and 32 bp, Tm = 71 °C ± 5.4 °C (the Tm difference between a primer pair was max. 3.9 °C), and a GC content of max. 61.5%. We used the previously published 515F/806R primers^52^ to amplify the 16S rRNA gene of the community in each sample. Standard curves for each strain were prepared in duplicates as a 10-fold dilution series, starting at a concentration of 10 ng template, with the lowest concentration being 0.1 pg template, to determine the linear range of each primer pair and the efficiency. All primer pairs demonstrated good linearity across at least 5 orders of magnitude (10 ng – 1 pg) (R^2^ of 0.9767 – 0.9995). Efficiencies for species-specific primers were between 70.5% and 94.3%, and efficiencies for the 16S rRNA gene primers were between 63.9% and 82.1%. The DNA of all cultures was extracted using the NucleoSpin Microbial DNA extraction kit (Macherey Nagel GmbH & Co. KG). The qPCR was performed using SYBR™ Green Universal Master Mix (Thermo Fisher Scientific). Each reaction contained 10 μL of the master mix, 7 μL nuclease-free H_2_O, 2 μL of 10 μM primer mix (final concentration 0.5 μM), and 1 μL template (4 ng – 10 ng). Samples were run in MicroAmp™ EnduraPlate™ Optical 96-Well Fast Clear Reaction Plates (Thermo Fisher Scientific) on a QuantStudio™ 1 Real-Time PCR System (Thermo Fisher Scientific), following the instructions in the SYBR™ Green Universal Master Mix manual (initial hold: 10 min at 95 °C, 40 cycles of denaturation [15 sec at 95 °C] and annealing/extension [1 min at 60 °C]). A melt curve analysis was performed for each sample followed by relative abundance analysis as described before ^53^. In brief, the ΔC_T_ value was determined for each species-specific gene in relation to the 16S rRNA gene of the co-culture, and the final OD_600_ of the culture was considered. With this method, we calculated the OD_600_ contribution of each strain to the final OD_600_ of the culture. The qPCR results were analyzed using the Design & Analysis 2.6.0 software (Thermo Fisher Scientific).

### Sample preparation for proteome analysis

To analyze the proteome of our single- and co-cultures, cells from 10 mL of each 24-hour culture were harvested by centrifugation (5 min, 15000 rcf, -9 °C), the supernatant was discarded, and the pellets were frozen at -70 °C overnight. The following day the cell pellets were resuspended in 1 mL of pre-cooled resuspension buffer (100 mM Tris-HCl pH 7.5 [Sigma-Aldrich, CAS 1185-53-1], 1 mM EDTA [Carl Roth, CAS 6381-92-6], 150 mM NaCl [Fisher Scientific, CAS 7647-14-5], 1 tablet of cOmplete™, Mini, EDTA-free Protease Inhibitor Cocktail [Roche, Cat. Nr. 04693159001], and 50 μL of 1 M DTT [Thermo Fisher Scientific, Cat. Nr.: R0861] per 10 mL buffer) and transferred into a Bead Tube Typ B (Macherey-Nagel, Cat. Nr.: 740812.50). Cells were disrupted using a FastPrep-24™ 5G bead-beating grinder and lysis system (MPbio), 4x 30 sec at 6.5 m/s. The samples were cooled on ice for 5 min between each bead-beating cycle. Subsequently, the lysis tubes were centrifuged for 20 min at 4 °C and 2000 rcf, and 200 μL of the supernatant was transferred to a new tube. The samples were stored at -70 °C until analysis. The proteome of biological triplicates was analyzed.

### Protein measurement

Proteins were precipitated with ice-cold acetone-methanol at -20 °C overnight. After centrifugation (2000 x g, 20 min, 4 °C), protein pellets were washed with 80 vol-% ice-cold acetone, and dried proteins were resolved in denaturation buffer (6 M urea, 2 M thiourea, 10 mM Tris, pH 8.0). Protein concentration was determined with Bradford assay, and 10 µg of proteins were subjected to in-solution digestion with trypsin: first, disulfide bonds were reduced with 1 mM DTT, and thiol groups of cysteine residues were carbamidomethylated with 5.5 mM iodoacetamide. Both steps were done at room temperature (RT) for 1 hour, in the dark. Then, a pre-digestion with endoproteinase LysC (Waco chemicals) in a peptidase-protein ratio of 1:100 weight-% was done at room temperature for 3 hours. After adding a fourfold volume of 20 mM ammonium bicarbonate to the peptide mixture, the final digestion with trypsin (Promega) was performed in a peptidase-peptide ratio of 1:100 weight-% overnight at room temperature. The reaction was stopped with trifluoroacetic acid (TFA) in a final concentration of 1 vol-%. Received peptides were desalted and stored on C18 StageTips^54^ until measurement.

LC-MS/MS analyses were performed on an EASY-nLC 1200 ultra-high performance liquid chromatography (UHPLC) system coupled to an Orbitrap Exploris 480 mass spectrometer through a nano-electrospray ion source (all Thermo Scientific) as described previously by Nashier, et al. (2024) with slight modifications: 0.25 µg peptides were loaded onto the analytical HPLC-column at a flow rate of 1.5 µL/min under a maximum back-pressure of 850 bar and subsequently eluted with a 46 min segmented gradient of 10-33-50 vol-% of HPLC solvent B at a flow rate of 200 nL/min.

### MS data processing and metaproteomics analysis

MS data from each co-culture and corresponding mono-cultures were processed together using default parameters of the MaxQuant software version 2.2.0.0^56^. Using the integrated Andromeda search engine^57^, obtained peak lists were searched against respective databases of the different organism, and 286 commonly observed contaminants. Peptide, protein, and modification-site identifications were reported at a false discovery rate (FDR) of 0.01, estimated by the target-decoy approach^58^. The intensity-based absolute quantification (iBAQ) and Label-Free Quantification (LFQ) algorithms^59^ were set, and “match between runs” was enabled.

To estimate the proportion of both bacteria in the co-culture and compare the expression of proteins between the monoculture and co-culture, we calculated a correction factor for each protein in each strain. Briefly, we calculated the sum of iBAQs (Σ iBAQs) for each sample that is on average highly correlated with protein abundance, followed by the calculation of the correction factor: CF = Average iBAQs co-culture/ Average iBAQs monoculture. For missing values in the co-culture CF = 1 and for missing values in the monocultures CF = to the CF obtained for each strain. Then each monoculture was analyzed versus its corresponding co-culture independently after protein annotation.

Downstream statistical analysis of proteomics data was performed in R (version 4.2.2). The R package Proteus (version 0.2.16) was used to analyze MaxQuant’s Proteomics output file “proteinGroups.txt”. Differential expression (DE) analysis was performed with the R package Limma (version 3.46.0) outside of the package Proteus. As a cut-off for statistical significance, an adjusted p value⍰(adj. P. Val.) ⍰< ⍰0.05 was chosen. All data were quantile normalized to account for the variation of intensity between samples followed by log2 transformation.

To identify differentially expressed proteins a linear model was then fitted to each protein as follows: exp⍰= ⍰ ⍰∼ ⍰condition with “exp” representing the expression of a protein and “condition” representing type of culture with the two levels monoculture and co-culture. For graphical visualization we used volcano plots to show statistical significance -log10(adj. *P*. Val) versus log2 fold change. The analysis script for one example co-culture experiment can be found on GitHub (https://github.com/qbic-projects/Q1130). For each analysis the specific ProteinGroups.txt of the commensal strain was used.

### Whole genome sequencing and genome alignment

The genome of *C. perfringens* C36 and the *C. perfringens* C36 *Δplc* deletion strain were sequenced using the Oxford Nanopore MinION sequencing device with an R9 and R10 flow cell, respectively. The DNA was prepared following the Rapid Barcoding Sequencing (SQK-RBK004) protocol and the Ligation sequencing DNA V14 (SQK-LSK14) protocol, respectively. The ONT tool Medaka v2.0.1 (https://github.com/nanoporetech/medaka?tab=readme-ov-file) was used to create consensus sequences from the sequencing data and the assembly was done with Flye v2.9.5-b1801^60^. For sequence annotation, we used Bakta^61^. Deletion validation was performed using GeneGone, a Python/Streamlit application developed in this study. GeneGone integrates Minimap2 (v2.28), BWA (v0.7.17), and samtools (v1.17) for read alignment and coverage profiling. Users provide a wild-type reference genome, sequencing reads, and the target locus. The tool calculates mean coverage across flanking regions and compares this to the target region. A deletion is called if coverage within the target locus falls below an adaptive threshold: by default, at least 20× coverage or 20% of flanking coverage for Illumina data, and at least 10× coverage or 20% of flanking coverage for long-read data. Although demonstrated here with *C. perfringens*, GeneGone is applicable to any organism where both sequencing data of a deletion strain and a wild-type reference genome are available. The software is openly available on GitHub (https://github.com/lucast122/GeneGone) and archived on Zenodo (DOI: 10.5281/zenodo.17076411). To further confirm that the gene deletion had taken place, the nucleotide sequence of *plc* was aligned against the corrected assembly of the previously sequenced *C. perfringens Δplc* genome using Minimap2 (v2.28-r1209)^62^ with the -x sr preset (optimized for short sequence alignment). As a control, the same gene sequence was also aligned to the wildtype genome using the same Minimap2 options, which revealed a perfect match at position 2,595,336 in the wildtype. To check for any additional deletions or larger genomic rearrangements, the full *C. perfringens* Δplc genome was aligned to the full wildtype genome using Minimap2 with the -x asm5 preset (suitable for similar genome assemblies). Inspection of the resulting alignments showed that the deleted gene had no alignment in the mutant genome, confirming its absence and that no other large-scale differences were detectable between the mutant and wildtype.

### Plasmid construction

An overview of all primers used for genetic modification of *C. perfringens* is given in **Supplementary Table 3**, and all plasmids are listed in **Supplementary Table 4**. All PCRs were performed with the Q5 Hot Start high-fidelity polymerase (NEB). *E. coli* NEB5α cells (NEB) were transformed with the ligation mix, and the plasmid was extracted from the strain and transferred into *E. coli* ST18^63^ for conjugation. The inserts of all plasmids were confirmed by Sanger sequencing (GENEWIZ Germany GmbH) of the insert region on the plasmids.

To generate the plasmid harboring the sgRNA and the repair template for the *plc* deletion (pMTL83251-*Δplc*), the *C. perfringens* P_*fdx*_ promoter region was amplified using primers CperfPfdx-fwd and Pfdx-plc-rev, and *C. perfringens* gDNA as template. The sgRNA spacer sequence was manually determined adjacent to a PAM sequence (NGG) and the sgRNA was amplified using the overlapping primers gRNA+term f2 and gRNA+term r2 without any template. Two 600-bp fragments were amplified up and downstream of *plc* using primers plc-RHA-f/plc-RHA-r and plc-LHA-f/plc-LHA-r, respectively. All primers introduced overlaps to the fragments, to the adjacent fragments, or to the vector. All inserts were first stepwise combined into one fragment by overlap-extension-PCR. pMTL83251^64^ was digested with KpnI (NEB) and NcoI (NEB) and the digested vector and the insert were assembled using the NEBuilder® HiFi DNA Assembly Master Mix (NEB). The plasmid for *iolE* deletion (pMTL83251-ΔiolE) was constructed following the workflow described for pMTL83251-*Δplc*, but the P_*fdx*_ promoter region was amplified with the reverse primer Pfdx-iolE-rev, primers gRNA-iolE-fwd and gRNA+term r2 were used to amplify the sgRNA, and for the repair template, we used primers LHA-iolE-fwd/LHA-iolE-rev and RHA-iolE-fwd/RHA-iolE-rev. To construct pMTL83151-P_*pfo*_-sLuc, harboring the luciferase gene under the control of the perfringolysin O promoter, the promoter region was amplified from *C. perfringens* gDNA using primers Ppfo-sLuc-fwd and LHA-pfo-rev. sLuc was amplified from pFF189^28^ using primers sLuc-pfo-fwd and Ppfo-sLuc-rev. pMTL83151^64^ was digested with NdeI (NEB) and NcoI (NEB) and the inserts assembled with the vector using the NEBuilder® HiFi DNA Assembly Master Mix (NEB).

### Plasmid transfer into C. perfringens

Plasmids were introduced into *C. perfringens via* conjugation with the *E. coli* ST18 mating strain^63^. The conjugation protocol was adapted from a previously described protocol from *C. difficile*^65^. The *E. coli* donor strain, carrying the plasmid to be transferred, and the recipient strain were inoculated in liquid medium and incubated 16 – 20 hours before being diluted 1:20 (*E. coli*) and 1:100 (*C. perfringens*) in fresh medium. Those cultures were incubated for at least 5 hours. Donor cells were harvested from 1 mL culture, washed once with 1X PBS and the pellet was transferred into the anaerobic chamber where it was resuspended in 25 μL – 50 μL *C. perfringens* culture. This cell suspension was spotted on a mGAM plate supplemented with 5-aminolevulinic acid and incubated overnight. The biomass from the mating spots was scraped off and transferred to a new plate supplemented with the selection antibiotics but omitting 5-aminolevulinic acid. Single colonies appeared after one or two days and were restreaked on new plates at least twice.

### Genetic modification of C. perfringens

To generate clean deletions in *C. perfringens*, we applied the RiboCas system^66^. Instead of including all components for the CRISPR/Cas9 system on one plasmid, we first introduced the Cas9 on pRGCas1 into *C. perfringens* and subsequently introduced a second plasmid harboring the sgRNA and the repair template.

The plasmids were transferred into *C. perfringens* via two consecutive conjugations as described above. The biomass from the second conjugal plasmid transfer was restreaked on a fresh mGAM plate supplemented with thiamphenicol, erythromycin, and theophylline to induce Cas9 expression. Colonies growing on this selection plate were tested for successful gene deletion *via* colony PCR (PhirePlant MasterMix), using biomass from one colony directly as template, and clones showing the gene deletion were transferred into liquid medium. The DNA was extracted, and the deletion region was sequenced by Sanger sequencing (GENEWIZ Germany GmbH) to confirm a clean deletion. For plasmid curing, the cultures were diluted three times in fresh medium at 42 °C and subsequently tested for antibiotic resistance and for plasmid presence *via* PCR with plasmid-specific primers.

### Perfringolysin O quantification

To quantify perfringolysin O in the supernatant of *C. perfringens* cultures, we established a sandwich enzyme-linked immunosorbent assay (ELISA). We followed the colorimetric sandwich ELISA protocol from ThermoFisher Scientific (https://assets.thermofisher.com/TFS-Assets/BID/Technical-Notes/general-elisa-protocols-tech-note.pdf). The plate was incubated overnight with 1 μg/mL Anti-*Clostridium perfringens* pfo/Perfringolysin O/Theta-toxin Antibody (HS3) (ProteoGenix). For the standard curve, we used Recombinant *Clostridium perfringens* pfo/Perfringolysin O/Theta-toxin Protein (ProteoGenix), and for antigen detection, we used 0.5 μg/mL primary detection antibody, *Clostridium perfringens* Perfringolysin O Polyclonal Antibody, PA5-117550 (Thermo Fisher Scientific) and 0.03125 μg/mL (1:32000) secondary antibody, Donkey anti-Rabbit-HRP, A16035 (Thermo Fisher Scientific). Upon incubation with TMB substrate solution (Thermo Fisher Scientific), the reaction was stopped by adding stop solution (Thermo Fisher Scientific), and the absorbance was measured in a plate reader (Tecan Infinite 200 PRO®) at 450 nm. All buffers and solutions required for the assay were purchased as ELISA Buffer Kit (CNB0011, Thermo Fisher Scientific). The protein concentration was determined using arigo’s ELISA Calculator, GainData® (https://www.arigobio.com/ELISA-calculator). Samples were analyzed using a four-parameter logistic (4PL) curve as a regression model. The OD_600_ contribution of *C. perfringens* in the co-cultures was determined by qPCR as described before and this value was used to calculate the protein concentration per OD_600_.

### Luminescence assay to determine the P_pfo_ promoter activity

The luminescence assay was performed as described before^29^. After 3 hours, 5 hours, 8 hours, 12 hours, and 24 hours of co-culture incubation, samples were taken for OD_600_ measurement, gDNA extraction and subsequent qPCR analysis, and for the luminescence assay. To measure the luminescence, each sample with OD_600_ > 0.3 was diluted 1:100, and 100 μL of the cell suspension was mixed with 20 μL of 1:50 diluted Nano-Glo® Luciferase Assay System substrate (Promega). After 15 minutes of incubation time, the luminescence was recorded in a plate reader (Tecan Infinite 200 PRO®). The OD_600_ of *C. perfringens* in the co-cultures was determined by qPCR as described before and this value was used to calculate the promoter activity (luminescence signal) per OD_600_. The background luminescence signal (signal from empty wells) was subtracted from the luminescence sample signals prior to normalization to the OD_600_.

### Sample preparation for targeted LC-MS/MS for amino acid measurement and metabolite profiling via FI-MS

1 mL of each single- and co-culture was centrifuged for 5 min at -9 °C and 18000 rcf. The supernatant was discarded, the cell pellets were dissolved in 1 mL quenching solution (40 vol-% acetonitrile [Sigma-Aldrich, CAS 75-05-8], 40 vol-% methanol [Sigma-Aldrich, CAS 67-56-1], 20 vol-% water), and the samples were kept at -20 °C overnight. The samples were then transferred to Bead Tubes Typ B (Macherey-Nagel, Cat. Nr.: 740812.50) and the cells were disrupted using a FastPrep-24™ 5G bead beating grinder and lysis system (MPbio), 2x 30 sec, 6.5 m/s. Subsequently, the samples were centrifuged 15 min at -9 °C and 13121 rcf and 400 μL of the supernatants were transferred into a fresh tube and stored at -70 °C until analysis. All steps were performed as quickly as possible and on ice.

### LC-MS/MS analysis for amino acid quantification

The analysis was performed on an Agilent TQ 6495 as described elsewhere^67^. The cell lysates were mixed 1:1 with the ^13^C internal standard and diluted 1:20 in quenching solution prior to measurement.

### FI-MS sample analysis

Samples were analyzed as described elsewhere with few adaptations^68^. An Agilent 6546 Series quadrupole time-of-flight mass spectrometer (Agilent Technologies, USA) was used for FI-MS and the electrospray source was operated in negative and positive ionization mode. For both ionization modes, the same mobile phase was used (60:40 vol-% of isopropanol:water, buffered with 10 mM ammonium carbonate and 0.04 vol-% ammonium hydroxide) at a flow rate of 0.15 mL/min. For online mass axis correction for the negative mode, we used 2-propanol (in the mobile phase) and HP-921, and for the positive mode purine and HP-921. Mass spectra were recorded in profile mode from 50 to 1,700 m/z with a frequency of 1.4 spectra/s for 0.5 min using 10 Ghz resolving power. The source temperature was 225 °C, with 11 L/min drying gas, and a nebulizer pressure of 20 psi. All other settings were kept as in the reference protocol. Raw Files (.d) were converted into .mzXML files using MSConvert (Settings: 64-bit, Write index, TPP compatibility, no peak-picking)^69^.

Data preprocessing was done as described before^68^. The metabolites were annotated using the Microbial Metabolites Database (MiMeDB)^70^. To account for the cell mass of each culture, the metabolite intensities were first normalized to the final OD_600_ of the culture and subsequently normalized to the mean intensity of each metabolite across all samples. The amino-acid intensities were extracted from the full data set and the PCA for each phylogenetic group was done with Metaboanalyst^71^. The heatmap for amino-acid intensities was created in R 4.4.1^72^ using the pheatmap package (Kolde R (2019). _pheatmap: Pretty Heatmaps_. R package version 1.0.12, https://CRAN.R-project.org/package=pheatmap).

### Sample preparation for non-targeted liquid chromatography-tandem mass spectrometry (LC-MS/MS)

#### Cell extract samples

Cells from 10 mL of each culture were harvested by centrifugation (5 min, -9 °C, 15000 rcf) and the cell pellet was resuspended in 1 mL quenching solution (40 vol-% acetonitrile (Sigma-Aldrich, CAS 75-05-8), 40 vol-% methanol (Sigma-Aldrich, CAS 67-56-1), 20 vol-% water). Samples were kept at -20 °C overnight before being lysed in Bead Tubes Typ B (Macherey-Nagel, Cat. Nr.: 740812.50) using a FastPrep-24™ 5G bead-beating grinder and lysis system (MPbio) 2x 30 sec, 6.5 m/s. Next, the lysates were centrifuged (15 min, -9 °C, 13121 rcf), and 400 μL of the supernatant was transferred to a 2 mL HPLC vial (Thermo Fisher Scientific, Cat. Nr.: 6PSV9-1P). The samples were stored at -70 °C until further processing.

#### Exometabolome samples

To extract the metabolites from the culture supernatant, 500 μL of supernatant from each culture used for cell extract metabolite analysis was transferred to a tube containing 1 mL precooled ethyl acetate (Sigma-Aldrich, CAS 141-78-6). The sample was vortexed twice for 30 sec and centrifuged for 5 min at -9 °C and 10000 rcf. 700 μL of the organic phase was carefully transferred to a 2 mL HPLC vial (Thermo Fisher Scientific, Cat. Nr.: 6PSV9-1P) and stored at -70 °C until it was further processed together with the cell-extract samples.

#### All sample types for LC-MS/MS

All samples for LC-MS/MS analysis were then dried down inside the HPLC vials using a speedvac. Each HPLC vial was weighed before samples were added, and the samples were weighed again after drying to determine the metabolite weight in each sample. The samples were resuspended in 50 vol-% methanol (LC/MS grade, FisherScientific) to a final concentration of 3 mg/mL and stored at -70 °C until analysis.

### Non-targeted LC-MS/MS analysis

Non-targeted metabolomics by LC-MS/MS was performed with an UHPLC coupled to a Q-Exactive HF mass spectrometer. In brief, the UHPLC separation was performed using a C18 core-shell column (Kinetex, 50 × 2.1 mm, 1.7 µm particle size, 100 A pore size, Phenomenex, Torrance, USA). The mobile phases used were solvent (A) water (LC/MS grade, Fisher Scientific) + 0.1 vol-% formic acid and solvent (B) acetonitrile (LC/MS grade, Fisher Scientific) + 0.1 vol-% formic acid. After the sample injection, a linear gradient method of 5 minutes was used for elution of small molecules, the flow-rate was set to 500 µL/min. The following separation conditions were set: in the time range 0-4 min from 5 vol-% to 50 vol-% solvent (B) was used, 4-5 min from 50 vol-% to 99 vol-% B, followed by 2 min washout phase at 99 vol-% B and 3 min re-equilibration phase at 5 vol-% B. The measurements were conducted in positive mode, the heated electrospray ionization (HESI) parameters included a sheath gas flow rate of 50 AU, auxiliary gas flow rate of 12 AU, and sweep gas flow rate of 1 AU. The spray voltage was set to 3.50 kV and the inlet capillary temperature to 250 °C, the S-lens RF level to 50 V and the auxiliary gas heater temperature to 400 °C. Full MS survey scan acquisition range was set to 150 –1,500 *m/z* with a resolution of 30,000, automatic gain control of 1·10^6^ (1E6), and maximum injection time of 100 ms with one micro-scan. MS/MS spectra acquisition was performed in data-dependent acquisition mode according to the optimized method from Stincone, *et al*. (2023). Briefly, TopN was set to 5, and consequently, the five most abundant precursor ions of the survey MS scan were destined to MS/MS fragmentation. The resolution of the MS/MS spectra was set to 15,000, the automatic gain control target to 5·10^5^ (1E6), and the maximum injection time to 50 ms. The quadrupole precursor selection width was set to 1 *m/z*. Normalized collision energy was applied stepwise at 25, 35, and 45. MS/MS scans were triggered with apex mode within 2 – 15 s from their first occurrence in a survey scan. Dynamic precursor exclusion was set to 5 s.The raw data files were converted to mzML format files using msConvert^74^ prior to being uploaded to MZmine 4.3.0^75^ for data processing. The output files, including feature tables and spectra information files, were used to create feature-based molecular networking (FBMN) in GNPS2^76^. SIRIUS output files generated by MZmine were used for metabolite class prediction with CANOPUS^37^, a tool integrated in SIRIUS 5.8.3^77^ (all settings are given in **Supplementary Table 5**). The link generated by GNPS2 was used to directly load the output into the FBMN-STATS app for statistical analysis (https://fbmn-statsguide.gnps2.org/)^78^. Pie charts were created with R 4.2.1^79^ using the ggforce package (Pedersen T (2024). ggforce: Accelerating ‘ggplot2’_. R package version 0.4.2, https://CRAN.R-project.org/package=ggforce). The heatmap for sphingolipid intensity was created in R 4.2.1^79^ using the pheatmap package (Kolde R (2019). _pheatmap: Pretty Heatmaps_. R package version 1.0.12, https://CRAN.R-project.org/package=pheatmap).

## Declarations

### Ethics approval and consent to participate

Not applicable

### Consent for publication

Not applicable

### Availability of data and materials

The 16S rRNA gene sequencing data supporting the conclusion of this article and the *C. perfringens* C36 and *C. perfringens* C36 Δplc genome sequences are available at the European Nucleotide Archive through accession IDs PRJEB76870 and PRJEB106409 (https://www.ebi.ac.uk/ena/browser/view/PRJEB76870 and https://www.ebi.ac.uk/ena/browser/view/PRJEB106409). The proteome dataset supporting the conclusions of this article is available at the ProteomeXchange Consortium through the PRIDE database^80,81^ with the project accession PXD074154, (https://www.ebi.ac.uk/pride/archive). The FI-MS dataset and raw and processed data files from the LC-MS/MS analyses were deposited to the Zenodo repository and can be found with the DOI 10.5281/zenodo.18269852 (https://doi.org/10.5281/zenodo.18269852).

### Competing interests

The authors declare that they have no competing interests

### Funding

This work was supported by the CMFI Cluster of Excellence in the framework of the Deutsche Forschungsgemeinschaft (DFG, German Research Foundation) under Germany’s Excellence Strategy – EXC 2124. Daniel Petras (DP) was supported by the National Institute of General Medical Sciences, GM160154, and Paolo Stincone (PS) was supported by the European Union’s Horizon Europe research and innovation programme through a Marie Skłodowska-Curie fellowship no. 101108450 MeStaLeM.

### Author’s contributions

Conceptualization of the project was carried out by Bastian Molitor (BM) and Lisa Maier (LM). Julia Schumacher (JS), LM, and BM designed the experiments. JS performed the genetic work, conducted the bioreactor experiments, the Oxford Nanopore MinION whole genome sequencing, the toxin quantification, and the assay to test the P_*pfo*_ promoter activity. JS and Carlos Llaca-Bautista (CLB) did the co-cultivation experiments, including qPCR analysis and sample preparation for proteome and metabolome analysis. Johanna Rapp (JR) and PS performed the metabolome sample analysis, and the proteome samples were analyzed by Mirita Franz-Wachtel (MFW) from the Proteome Center Tübingen. The statistical analysis of the proteome data was done by Francesca Barletta-Solari (FBS) from the Quantitative Biology Center (QBiC) and JS analyzed the metabolome data and pre-processed proteome data under the supervision of FBS, PS, JR, DP, Hannes Link (HL), and BM. Timo-Niklas Lucas (TNL) performed the genome assembly and annotation and analyzed the genome of the *Δplc* deletion strain under the supervision of Daniel H. Huson (DHH). LM provided the community strains and the plate reader for luminescence measurements. JS prepared all figures and tables. JS drafted the manuscript, and all authors contributed to editing the draft.

## Acknowledgements

We thank Dr. Libera Lo Presti for writing support and PD Dr. Meina Neumann-Schaal for valuable scientific discussions. We further thank Silke Wahl for the great support with protein sample preparation for mass spectrometry analysis.

## Additional files 1-2

### Additional file 1

File format: Microsoft Word-Document (.docx)

Title: Supplementary Figures

Description: contains Supplementary Figures 1-7, the corresponding figure legends, and references

### Additional file 2

File format: Microsoft Excel-Sheet (.xlsx)

Title: Supplementary Tables

Description: contains Supplementary Tables 1-5. Tables 1-4 contain information about strains, primers, and plasmids used in this study, Table 5 contains the settings used for LC-MS/MS data analysis with SIRIUS.

## Notes

### Competing Interest Statement

The authors have declared no competing interest.

## References

1 Freedman, J. C. et al. Clostridium perfringens type A-E toxin plasmids. Res Microbiol 166, 264–279, doi:10.1016/j.resmic.2014.09.004 (2015).

2 Rood, J. I. et al. Expansion of the Clostridium perfringens toxin-based typing scheme. Anaerobe 53, 5–10, doi:10.1016/j.anaerobe.2018.04.011 (2018).

3 Camargo, A., Ramírez, J. D., Kiu, R., Hall, L. J. & Muñoz, M. Unveiling the pathogenic mechanisms of Clostridium perfringens toxins and virulence factors. Emerging Microbes & Infections 13, 2341968, doi:10.1080/22221751.2024.2341968 (2024).

4 Labbe, R. G. & Huang, T. H. Generation times and modeling of enterotoxin-positive and enterotoxin-negative strains of Clostridium perfringens in laboratory media and ground beef. J Food Prot 58, 1303–1306, doi:10.4315/0362-028X-58.12.1303 (1995).

5 Shimizu, T. et al. Complete genome sequence of Clostridium perfringens, an anaerobic flesh-eater. Proc Natl Acad Sci U S A 99, 996–1001, doi:10.1073/pnas.022493799 (2002).

6 Goldner, S. B., Solberg, M. & Post, L. S. Development of a minimal medium for Clostridium perfringens by using an anaerobic chemostat. Applied and Environmental Microbiology 50, 202–206, doi:10.1128/aem.50.2.202-206.1985 (1985).

7 Fuchs, A.-R. & Bonde, G. J. The nutritional requirements of Clostridium perfringens. Microbiology 16, 317–329, doi:10.1099/00221287-16-2-317 (1957).

8 Grenda, T. et al. Clostridium perfringens-opportunistic foodborne pathogen, its diversity and epidemiological significance. Pathogens 12, doi:10.3390/pathogens12060768 (2023).

9 Yao, P. Y. & Annamaraju, P. Clostridium perfringens infection. StatPearls (2024).

10 Chen, Y., Xiao, L., Zhou, M. & Zhang, H. The microbiota: a crucial mediator in gut homeostasis and colonization resistance. Front Microbiol 15, 1417864, doi:10.3389/fmicb.2024.1417864 (2024).

11 García-Vela, S., Martínez-Sancho, A., Said, L. B., Torres, C. & Fliss, I. Pathogenicity and antibiotic resistance diversity in Clostridium perfringens isolates from poultry affected by necrotic enteritis in Canada. Pathogens 12, doi:10.3390/pathogens12070905 (2023).

12 Song, X. et al. Emergence of genetic diversity and multi-drug resistant Clostridium perfringens from wild birds. BMC Vet Res 20, 300, doi:10.1186/s12917-024-04168-8 (2024).

13 Khandia, R. et al. Wound infection with multi-drug resistant Clostridium perfringens: a case study. Arch Razi Inst 76, 1565–1573, doi:10.22092/ari.2021.355985.1757 (2021).

14 Zhong, J. x. et al. Molecular characteristics and phylogenetic analysis of Clostridium perfringens from different regions in China, from 2013 to 2021. Frontiers in Microbiology 14, doi:10.3389/fmicb.2023.1195083 (2023).

15 Geremia, N. et al. A subanalysis of Clostridium perfringens bloodstream infections from a 5-year retrospective nationwide survey (ITANAEROBY). Anaerobe 90, 102901, doi:10.1016/j.anaerobe.2024.102901 (2024).

16 Maier, L. et al. Unravelling the collateral damage of antibiotics on gut bacteria. Nature 599, 120–124, doi:10.1038/s41586-021-03986-2 (2021).

17 Yan, Z. et al. High intestinal carriage of Clostridium perfringens in healthy individuals and ICU patients in Hangzhou, China. Microbiol Spectr 12, e0338523, doi:10.1128/spectrum.03385-23 (2024).

18 Cree, B. A., Spencer, C. M., Varrin-Doyer, M., Baranzini, S. E. & Zamvil, S. S. Gut microbiome analysis in neuromyelitis optica reveals overabundance of Clostridium perfringens. Ann Neurol 80, 443–447, doi:10.1002/ana.24718 (2016).

19 Nagpal, R. et al. Sensitive quantification of Clostridium perfringens in human feces by quantitative real-time PCR targeting alpha-toxin and enterotoxin genes. BMC Microbiol 15, 219, doi:10.1186/s12866-015-0561-y (2015).

20 Hou, K. et al. Microbiota in health and diseases. Signal Transduct Target Ther 7, 135, doi:10.1038/s41392-022-00974-4 (2022).

21 Khan, I. et al. Mechanism of the gut microbiota colonization resistance and enteric pathogen infection. Front Cell Infect Microbiol 11, 716299, doi:10.3389/fcimb.2021.716299 (2021).

22 Martin, A. J. M., Serebrinsky-Duek, K., Riquelme, E., Saa, P. A. & Garrido, D. Microbial interactions and the homeostasis of the gut microbiome: the role of Bifidobacterium. Microbiome Res Rep 2, 17, doi:10.20517/mrr.2023.10 (2023).

23 Liu, Y. et al. Analysis of the dynamic changes in gut microbiota in patients with different severity in sepsis. BMC Infect Dis 23, 614, doi:10.1186/s12879-023-08608-y (2023).

24 Kullberg, R. F. J. et al. Rectal bacteriome and virome signatures and clinical outcomes in community-acquired pneumonia: An exploratory study. EClinicalMedicine 39, 101074, doi:10.1016/j.eclinm.2021.101074 (2021).

25 Singh, P. et al. Intestinal microbial communities associated with acute enteric infections and disease recovery. Microbiome 3, 45, doi:10.1186/s40168-015-0109-2 (2015).

26 Daquigan, N., Seekatz, A. M., Greathouse, K. L., Young, V. B. & White, J. R. High-resolution profiling of the gut microbiome reveals the extent of Clostridium difficile burden. NPJ Biofilms Microbiomes 3, 35, doi:10.1038/s41522-017-0043-0 (2017).

27 Yang, Q. et al. Identification of an intestinal microbiota signature associated with the severity of necrotic enteritis. Frontiers in Microbiology 12, doi:10.3389/fmicb.2021.703693 (2021).

28 Schumacher, J. et al. Proton-pump inhibitors increase C. difficile infection risk by altering pH rather than by affecting the gut microbiome based on a bioreactor model. Gut Microbes 17, 2519697, doi:10.1080/19490976.2025.2519697 (2025).

29 Grießhammer, A. et al. Non-antibiotic drugs break colonization resistance against pathogenic Gammaproteobacteria. bioRxiv, 2023.2011.2006.564936, doi:10.1101/2023.11.06.564936 (2023).

30 Galperin, M. Y. et al. COG database update: focus on microbial diversity, model organisms, and widespread pathogens. Nucleic Acids Res 49, D274–D281, doi:10.1093/nar/gkaa1018 (2021).

31 Pols, T., Singh, S., Deelman-Driessen, C., Gaastra, B. F. & Poolman, B. Enzymology of the pathway for ATP production by arginine breakdown. The FEBS Journal 288, 293–309, doi:10.1111/febs.15337 (2021).

32 Medlock, G. L. et al. Inferring metabolic mechanisms of interaction within a defined gut microbiota. Cell Syst 7, 245–257 e247, doi:10.1016/j.cels.2018.08.003 (2018).

33 Heaver, S. L. et al. Characterization of inositol lipid metabolism in gut-associated Bacteroidetes. Nat Microbiol 7, 986–1000, doi:10.1038/s41564-022-01152-6 (2022).

34 Kawsar, H. I., Ohtani, K., Okumura, K., Hayashi, H. & Shimizu, T. Organization and transcriptional regulation of myo-inositol operon in Clostridium perfringens. FEMS Microbiology Letters 235, 289–295, doi:10.1111/j.1574-6968.2004.tb09601.x (2004).

35 Urbina, P. et al. Unexpected wide substrate specificity of C. perfringens alpha-toxin phospholipase C. Biochim Biophys Acta 1808, 2618–2627, doi:10.1016/j.bbamem.2011.06.008 (2011).

36 Verherstraeten, S. et al. Perfringolysin O: The underrated Clostridium perfringens toxin? Toxins (Basel) 7, 1702–1721, doi:10.3390/toxins7051702 (2015).

37 Duhrkop, K. et al. Systematic classification of unknown metabolites using high-resolution fragmentation mass spectra. Nat Biotechnol 39, 462–471, doi:10.1038/s41587-020-0740-8 (2021).

38 Heaver, S. L., Johnson, E. L. & Ley, R. E. Sphingolipids in host-microbial interactions. Curr Opin Microbiol 43, 92–99, doi:10.1016/j.mib.2017.12.011 (2018).

39 Nguyen, T. H. N., Szamosvári, D. & Böttcher, T. Synthesis of bacterial 2-alkyl-4(1H)-quinolone derivatives. ARKIVOC 2021, 218–239, doi:10.24820/ark.5550190.p011.547 (2021).

40 Rossjohn, J. et al. Structures of perfringolysin O suggest a pathway for activation of cholesterol-dependent cytolysins. J Mol Biol 367, 1227–1236, doi:10.1016/j.jmb.2007.01.042 (2007).

41 Berman, H. M. et al. The Protein Data Bank. Nucleic Acids Research 28, 235–242, doi:10.1093/nar/28.1.235 (2000).

42 Rossjohn, J., Parker, M., Polekhina, G., Feil, S. & Tweten, R. in Worldwide Protein Data Bank (2004).

43 Weiss, A. S. et al. In vitro interaction network of a synthetic gut bacterial community. The ISME Journal 16, 1095–1109, doi:10.1038/s41396-021-01153-z (2022).

44 Wintermute, E. H. & Silver, P. A. Emergent cooperation in microbial metabolism. Molecular Systems Biology 6, 407, doi:10.1038/msb.2010.66 (2010).

45 Hong, Y.-J., Cai, Y. & Antoniewicz, M. R. Cross-feeding of amino acid pathway intermediates is common in co-cultures of auxotrophic Escherichia coli. Metabolic Engineering 88, 172–179, doi:10.1016/j.ymben.2025.01.003 (2025).

46 Ohtani, K. et al. Identification of a two-component VirR/VirS regulon in Clostridium perfringens. Anaerobe 16, 258–264, doi:10.1016/j.anaerobe.2009.10.003 (2010).

47 Ba-Thein, W. et al. The virR/virS locus regulates the transcription of genes encoding extracellular toxin production in Clostridium perfringens. J Bacteriol 178, 2514–2520, doi:10.1128/jb.178.9.2514-2520.1996 (1996).

48 Johnson, E. L. et al. Sphingolipids produced by gut bacteria enter host metabolic pathways impacting ceramide levels. Nat Commun 11, 2471, doi:10.1038/s41467-020-16274-w (2020).

49 Brown, E. M. et al. Bacteroides-derived sphingolipids are critical for maintaining intestinal homeostasis and symbiosis. Cell Host Microbe 25, 668–680.e667, doi:10.1016/j.chom.2019.04.002 (2019).

50 Ryan, E., Joyce, S. A. & Clarke, D. J. Membrane lipids from gut microbiome-associated bacteria as structural and signalling molecules. Microbiology (Reading) 169, doi:10.1099/mic.0.001315 (2023).

51 Truong, D. T. et al. MetaPhlAn2 for enhanced metagenomic taxonomic profiling. Nat Methods 12, 902–903, doi:10.1038/nmeth.3589 (2015).

52 Caporaso, J. G. et al. Global patterns of 16S rRNA diversity at a depth of millions of sequences per sample. Proc Natl Acad Sci U S A 108 Suppl 1, 4516–4522, doi:10.1073/pnas.1000080107 (2011).

53 Garcia-Santamarina, S. et al. Emergence of community behaviors in the gut microbiota upon drug treatment. Cell 187, 6346–6357 e6320, doi:10.1016/j.cell.2024.08.037 (2024).

54 Rappsilber, J., Mann, M. & Ishihama, Y. Protocol for micro-purification, enrichment, pre-fractionation and storage of peptides for proteomics using StageTips. Nature Protocols 2, 1896–1906, doi:10.1038/nprot.2007.261 (2007).

55 Nashier, P. et al. Deep phosphoproteomics of Klebsiella pneumoniae reveals HipA-mediated tolerance to ciprofloxacin. PLoS Pathog 20, e1012759, doi:10.1371/journal.ppat.1012759 (2024).

56 Cox, J. & Mann, M. MaxQuant enables high peptide identification rates, individualized p.p.b.-range mass accuracies and proteome-wide protein quantification. Nat Biotechnol 26, 1367–1372, doi:10.1038/nbt.1511 (2008).

57 Cox, J. et al. Andromeda: a peptide search engine integrated into the MaxQuant environment. J Proteome Res 10, 1794–1805, doi:10.1021/pr101065j (2011).

58 Elias, J. E. & Gygi, S. P. Target-decoy search strategy for increased confidence in large-scale protein identifications by mass spectrometry. Nat Methods 4, 207–214, doi:10.1038/nmeth1019 (2007).

59 Tyanova, S., Temu, T. & Cox, J. The MaxQuant computational platform for mass spectrometry-based shotgun proteomics. Nat Protoc 11, 2301–2319, doi:10.1038/nprot.2016.136 (2016).

60 Kolmogorov, M., Yuan, J., Lin, Y. & Pevzner, P. A. Assembly of long, error-prone reads using repeat graphs. Nature Biotechnology 37, 540–546, doi:10.1038/s41587-019-0072-8 (2019).

61 Schwengers, O. et al. Bakta: rapid and standardized annotation of bacterial genomes via alignment-free sequence identification. Microb Genom 7, doi:10.1099/mgen.0.000685 (2021).

62 Li, H. Minimap2: pairwise alignment for nucleotide sequences. Bioinformatics 34, 3094–3100, doi:10.1093/bioinformatics/bty191 (2018).

63 Thoma, S. & Schobert, M. An improved Escherichia coli donor strain for diparental mating. FEMS Microbiol Lett 294, 127–132, doi:10.1111/j.1574-6968.2009.01556.x (2009).

64 Heap, J. T., Pennington, O. J., Cartman, S. T. & Minton, N. P. A modular system for Clostridium shuttle plasmids. J Microbiol Methods 78, 79–85, doi:10.1016/j.mimet.2009.05.004 (2009).

65 Bouillaut, L., McBride, S. M. & Sorg, J. A. Genetic manipulation of Clostridium difficile. Curr Protoc Microbiol Chapter 9, Unit 9A.2, doi:10.1002/9780471729259.mc09a02s20 (2011).

66 Canadas, I. C., Groothuis, D., Zygouropoulou, M., Rodrigues, R. & Minton, N. P. RiboCas: a universal CRISPR-based editing tool for Clostridium. ACS Synth Biol 8, 1379–1390, doi:10.1021/acssynbio.9b00075 (2019).

67 Guder, J. C., Schramm, T., Sander, T. & Link, H. Time-optimized isotope ratio LC-MS/MS for high-throughput quantification of primary metabolites. Anal Chem 89, 1624–1631, doi:10.1021/acs.analchem.6b03731 (2017).

68 Farke, N., Schramm, T., Verhülsdonk, A., Rapp, J. & Link, H. Systematic analysis of in-source modifications of primary metabolites during flow-injection time-of-flight mass spectrometry. Analytical Biochemistry 664, 115036, doi:10.1016/j.ab.2023.115036 (2023).

69 Chambers, M. C. et al. A cross-platform toolkit for mass spectrometry and proteomics. Nat Biotechnol 30, 918–920, doi:10.1038/nbt.2377 (2012).

70 Wishart, D. S. et al. MiMeDB: the Human Microbial Metabolome Database. Nucleic Acids Res 51, D611–D620, doi:10.1093/nar/gkac868 (2023).

71 Pang, Z. et al. MetaboAnalystR 4.0: a unified LC-MS workflow for global metabolomics. Nat Commun 15, 3675, doi:10.1038/s41467-024-48009-6 (2024).

72 Team, R. C. (R Foundation for Statistical Computing, Vienna, Austria, 2024).

73 Stincone, P. et al. Evaluation of data-dependent MS/MS acquisition parameters for non-targeted metabolomics and molecular networking of environmental samples: Focus on the Q Exactive Platform. Anal Chem 95, 12673–12682, doi:10.1021/acs.analchem.3c01202 (2023).

74 Adusumilli, R. & Mallick, P. Data conversion with ProteoWizard msConvert. Proteomics: Methods and Protocols, 339–368, doi:10.1007/978-1-4939-6747-6_23 (2017).

75 Schmid, R. et al. Integrative analysis of multimodal mass spectrometry data in MZmine 3. Nat Biotechnol 41, 447–449, doi:10.1038/s41587-023-01690-2 (2023).

76 Wang, M. et al. Sharing and community curation of mass spectrometry data with Global Natural Products Social Molecular Networking. Nat Biotechnol 34, 828–837, doi:10.1038/nbt.3597 (2016).

77 Duhrkop, K. et al. SIRIUS 4: a rapid tool for turning tandem mass spectra into metabolite structure information. Nat Methods 16, 299–302, doi:10.1038/s41592-019-0344-8 (2019).

78 Pakkir Shah, A. K. et al. The Hitchhiker’s guide to statistical analysis of feature-based molecular networks from non-targeted metabolomics data. ChemRxiv, doi:10.26434/chemrxiv-2023-wwbt0 (2023).

79 R: A language and environment for statistical computing. (R Foundation for Statistical Computing, Vienna, Austria, 2022).

80 Deutsch, E. W. et al. The ProteomeXchange consortium at 10 years: 2023 update. Nucleic Acids Research 51, D1539–D1548, doi:10.1093/nar/gkac1040 (2022).

81 Perez-Riverol, Y. et al. The PRIDE database at 20 years: 2025 update. Nucleic Acids Res 53, D543–D553, doi:10.1093/nar/gkae1011 (2025).

