## Supplementary Figures for "A host-directed virulence factor of *Clostridium perfringens* is modulated by gut commensal strains"

Julia Schumacher^1,2^	ORCID0009-0001-4843-1075

Paolo Stincone^2^	ORCID0000-0002-2214-6655

Johanna Rapp^2,3^	ORCID0000-0002-4591-8312

Timo-Niklas Lucas^2,4^	ORCID

Carlos Llaca-Bautista^1^	ORCID0009-0004-2362-6986

Francesca Barletta^5,6^	ORCID0009-0002-1396-2252

Mirita Franz-Wachtel^7^	ORCID

Boris Maček^7^	ORCID0000-0002-1206-2458

Daniel H. Huson^8^	ORCID0000-0002-2961-604X

Lisa Maier^2,6,9^	ORCID0000-0002-6473-4762

Hannes Link^2,3^	ORCID0000-0002-6677-555X

Daniel Petras^2,10^	ORCID0000-0002-6561-3022

Bastian Molitor^1,2,11,^*	ORCID0000-0002-0776-1668

^1^ Environmental Biotechnology Group, Department of Geosciences, University of Tübingen, Germany

^2^ Cluster of Excellence EXC 2124: Controlling Microbes to Fight Infection (CMFI), University of Tübingen, Germany

^3^ Bacterial Metabolomics, Interfaculty Institute of Microbiology and Infection Medicine, University of Tübingen, Germany

^4^ Institute for Bioinformatics and Medical Informatics, University of Tübingen, Germany

^5^ Quantitative Biology Center (QBiC), University of Tübingen, Germany

^6^ M3-Research Center for Malignome, Metabolome and Microbiome, University Hospital, Germany

^7^ Proteome Center Tübingen, Institute of Cell Biology, University of Tübingen, Germany

^8^ Institute for Bioinformatics and Medical Informatics, University of Tübingen, Germany

^9^ Interfaculty Institute for Microbiology and Infection Medicine Tübingen, University of Tübingen, Germany

^10^ Department of Biochemistry, University of California Riverside, USA

^11^ Microbial Metabolic Biochemistry, Institute of Biochemistry, Leipzig University, Germany

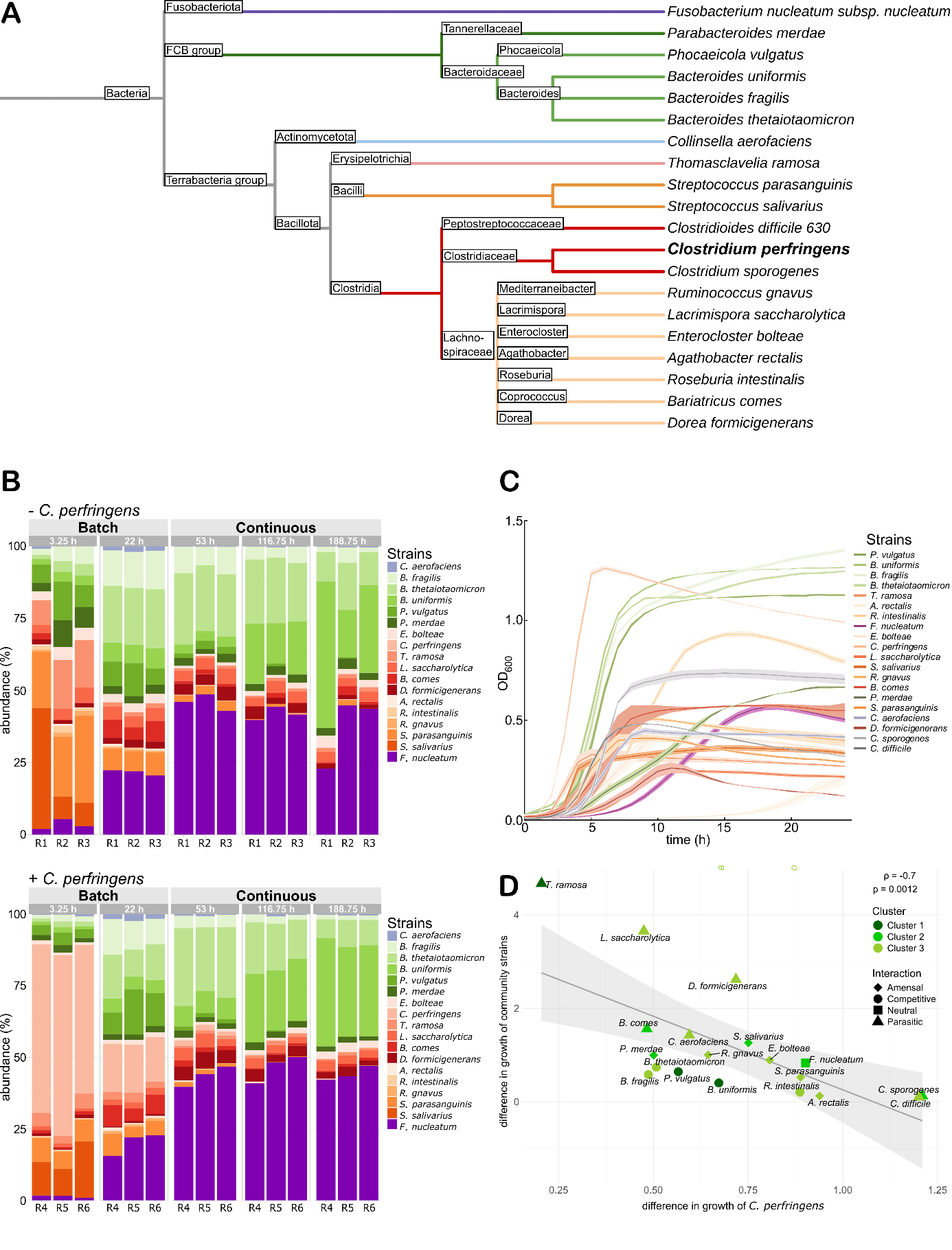

**Supplementary Figure 1: Com18 composition and growth of axenic cultures and co-cultures. A:** Phylogenetic tree of Com18 including *C. perfringens* and the two other pathobionts, *C. difficile* and *C. sporogenes*. The phylogenetic tree was generated using iTOL^33^. **B:** Relative abundance of each strain of Com18 (and *C. perfringens*) in the absence (top, “- *Clostridium perfringens*”) or presence (bottom, “+ *Clostridium perfringens*”) of *C. perfringens*. R1-3: bioreactor replicates without *C. perfringens*, R4-6: bioreactor replicates with *C. perfringens*, the triplicates are represented in one panel per timepoint. The first two time points originate from batch operation, and the following three time points from continuous operation. The relative abundances were determined *via* 16S rRNA sequencing. **C:** Growth curves of all strains in single cultures over 24 hours. The standard deviations are shown as areas adjacent to the mean growth (N = 3). **D:** The relative growth of *C. perfringens* in co-culture plotted against the relative growth of the community strains in co-culture. The grey line represents the linear trend line. The Spearman coefficient (ρ) and p-value are indicated in the top right. Each co-culture was colored based on the cluster into which the community strain falls as shown in **Figure 2A**, and the shape represents the interaction type.

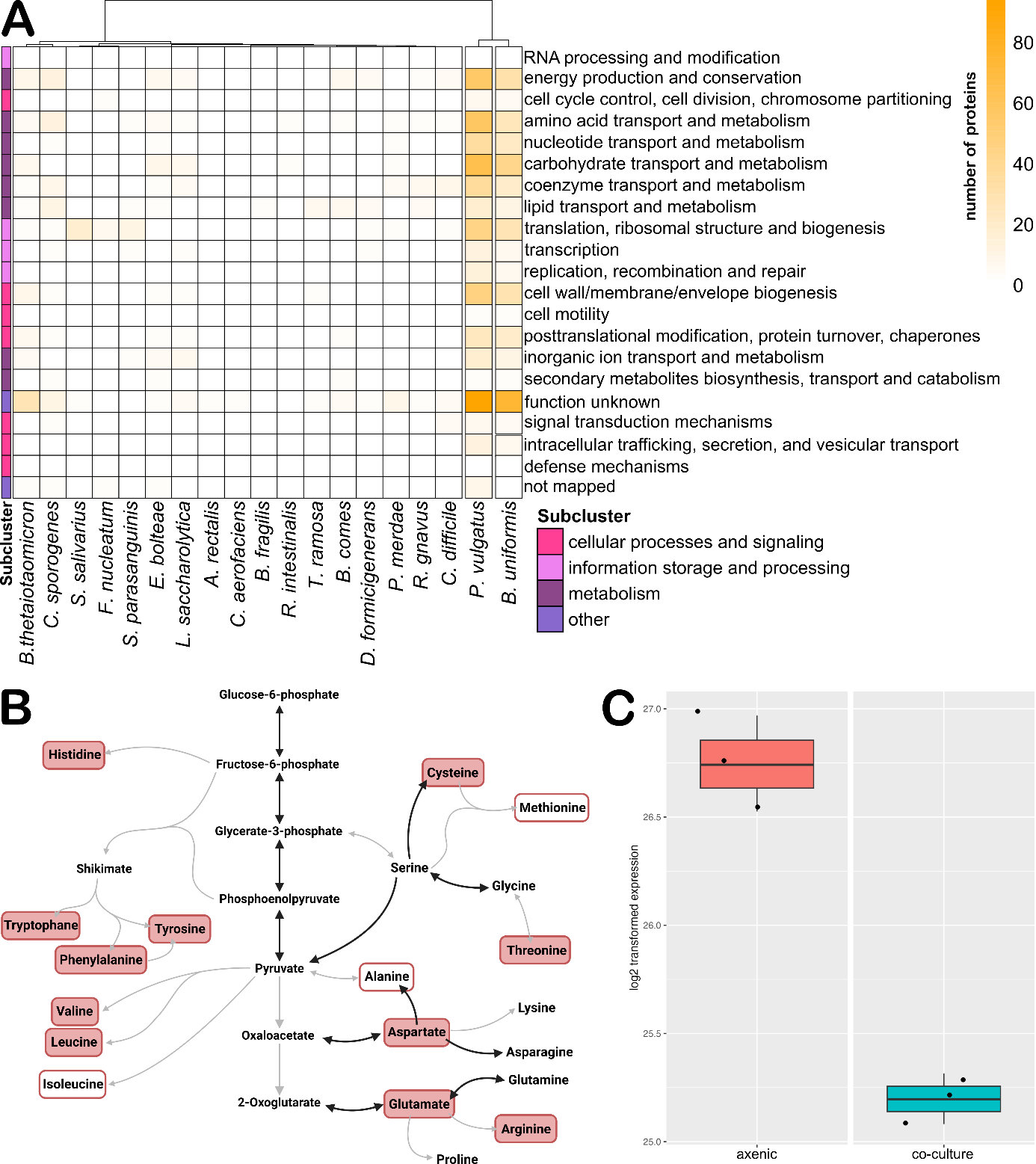

**Supplementary Figure 2: Competition for amino-acids in co-cultures. A:** Heatmap of proteins that were found to be more abundant in the co-cultures compared to the single culture. Each protein was classified according to the COG database and the number of proteins in each class is visualized in the heatmap. The strains are arranged according to the Euclidean distance metric. Mean values of triplicates were used. **B:** Amino acid biosynthesis pathway as shown by KEGG^1^ (map01230). *C. perfringens* enzymes annotated in the genome are shown as black arrows, and grey arrows symbolize absent genes. Amino acids in filled red boxes were shown to be essential in several studies. Amino acids with a red border were found to be essential in at least one publication^2-4^. **C:** Abundance of the *B. fragilis* protein Q5LEZ7 in the *B. fragilis* axenic culture (axenic) and in the *C. perfringens* co-culture (co-culture).

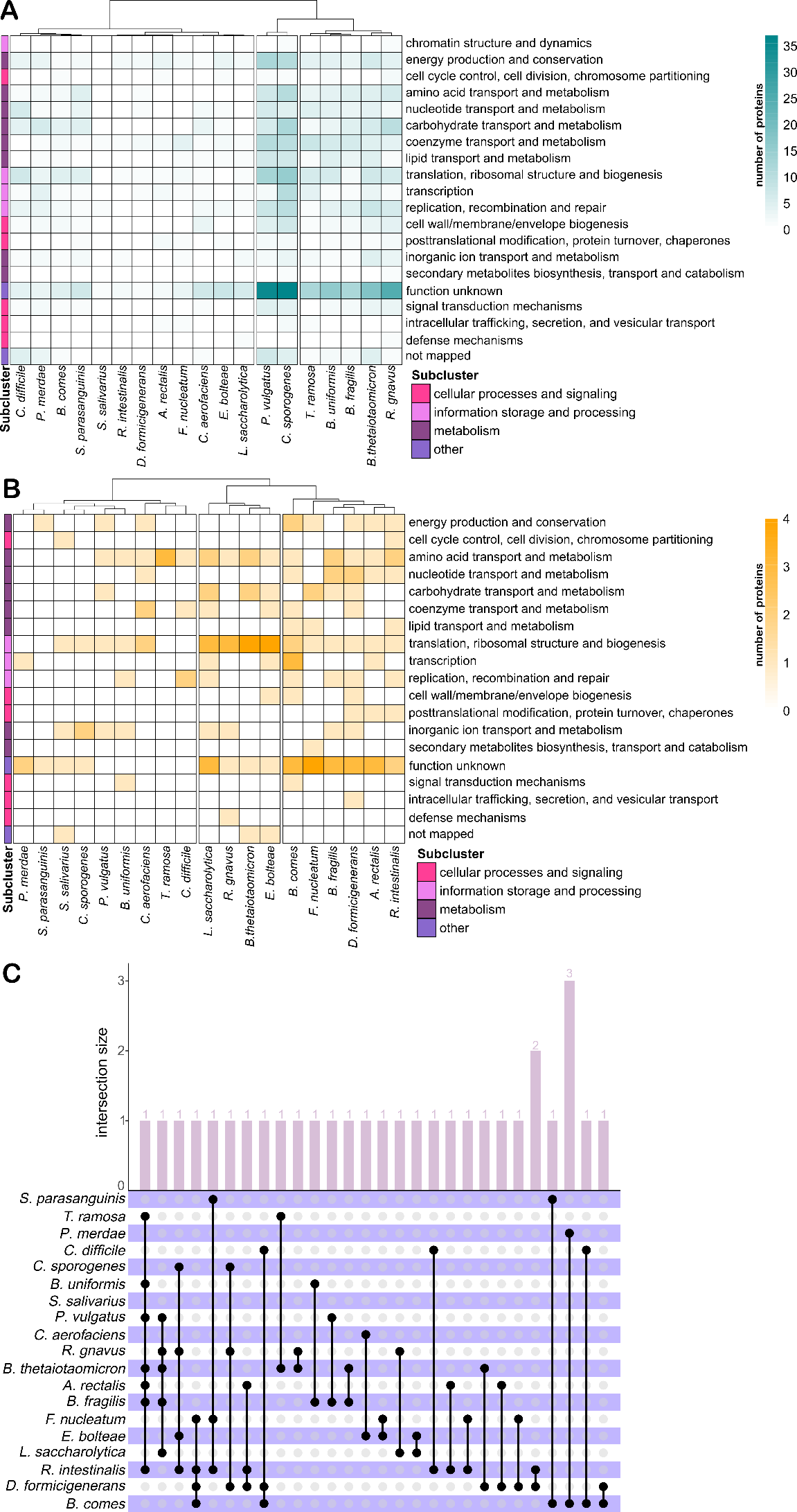

**Supplementary Figure 3: Proteome response of *C. perfringens* in co-cultures. A:** Heatmap of *C. perfringens* proteins that were found to be less abundant or absent in the co-cultures compared to the single culture. **B:** Heatmap of *C. perfringens* proteins that were found to be more abundant or present in the co-cultures compared to the single culture. **A and B:** Each protein was classified according to the COG database and the number of proteins in each class is visualized in the heatmap. Euclidean distance metric was used to determine the relationship. Mean values of triplicates were used. **C:** UpSet plot of *C. perfringens* proteins that are present or more abundant in the co-culture than the single culture. Only proteins that were found in at least two cultures are included.

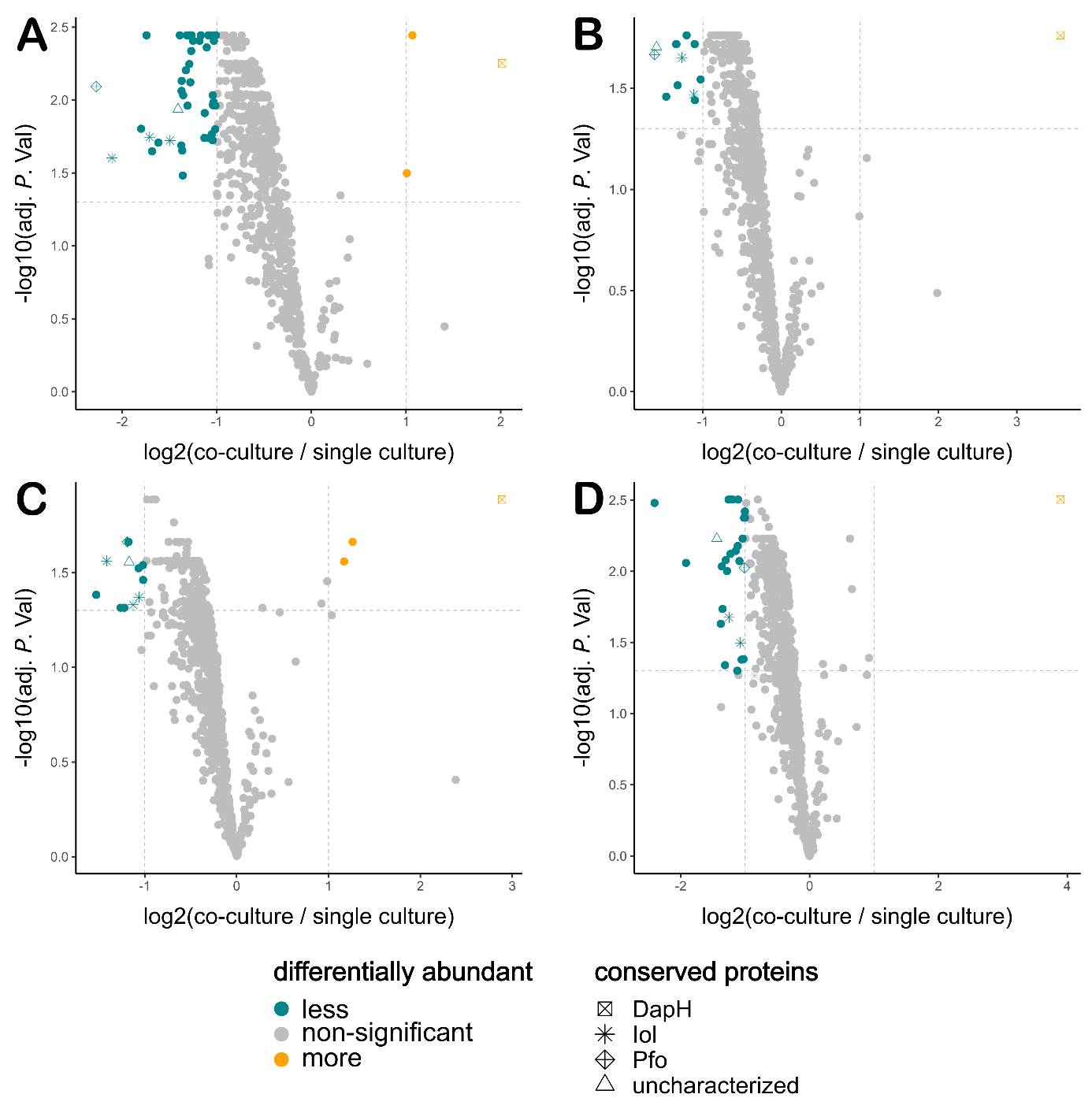

**Supplementary Figure 4:** **Differentially abundant *C. perfringens* proteins in co-cultures. With A:** *P. vulgatus***, B:** *B. uniformis***, C:** *B. fragilis*, and **D:** *B. thetaiotaomicron*. Proteins were considered to be differentially abundant if the fold change (co-culture / single culture) was above 2 (log2(co-culture / single culture) ≥ |2|) and the adjusted *p*-value ≤ 0.05 (-log10(adj. *P*. Val) ≥ 1.3). Mean values from replicates are shown (n=3).

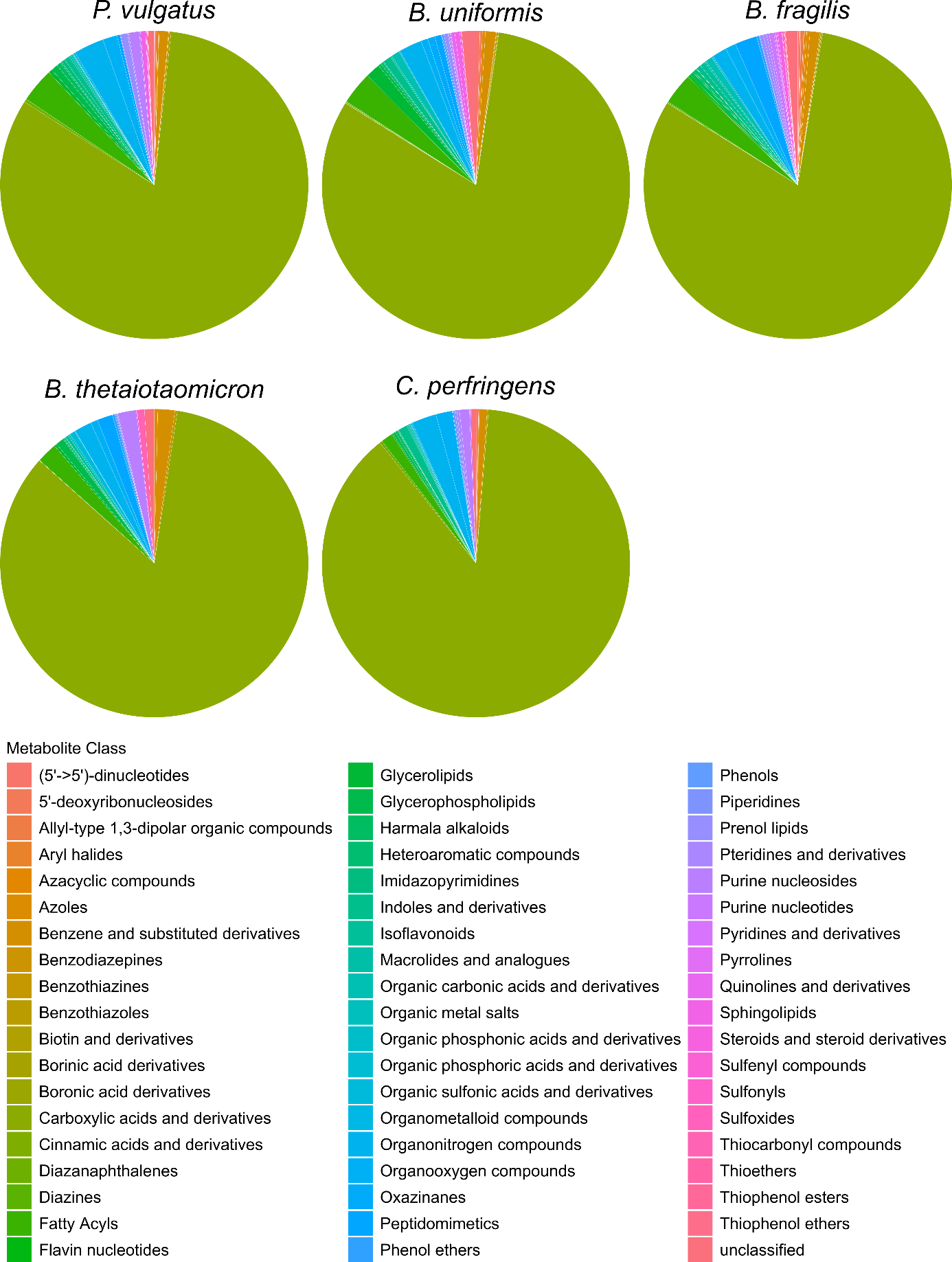

**Supplementary Figure 5: Metabolite classes of the cell-associated metabolomes.** Pie charts with metabolite classes found in the metabolomes of *P. vulgatus*, *B. uniformis*, *B. fragilis*, *B. thetaiotaomicron*, and *C. perfringens.*

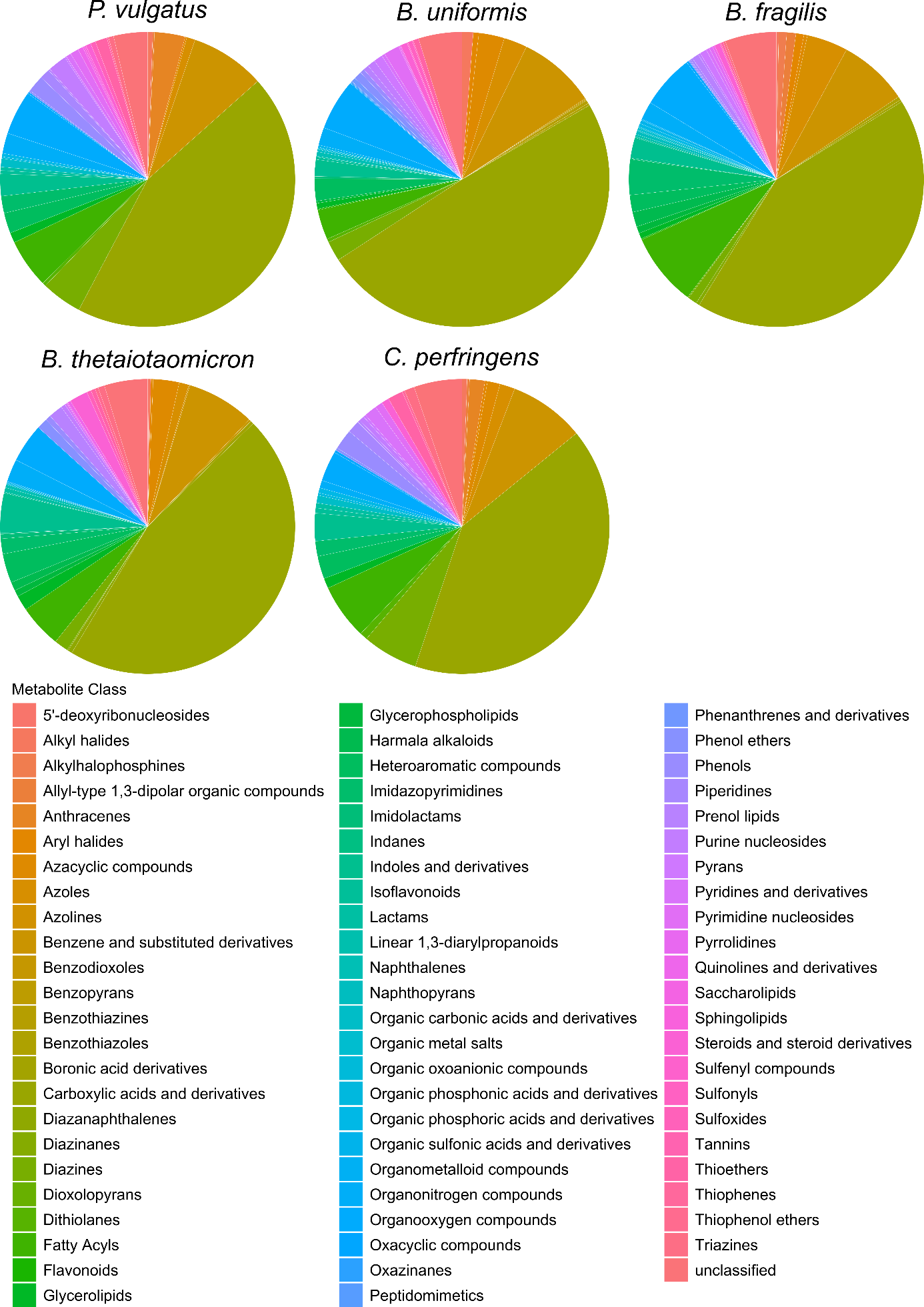

**Supplementary Figure 6: Metabolite classes of the exometabolomes.** Pie charts with metabolite classes found in the metabolomes of *P. vulgatus*, *B. uniformis*, *B. fragilis*, *B. thetaiotaomicron*, and *C. perfringens*.

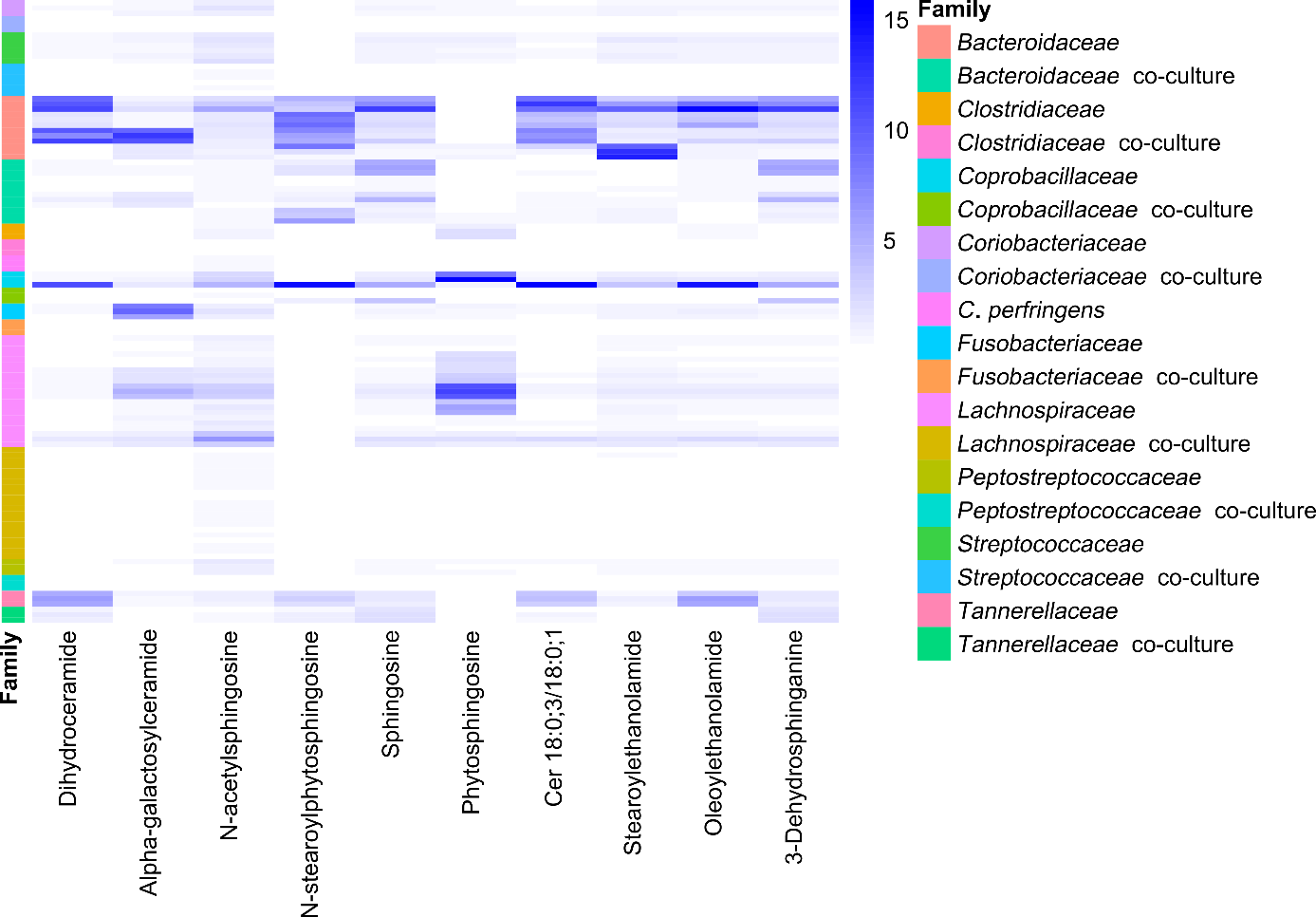

**Supplementary Figure 7: Sphingolipids in Com18 axenic cultures and co-cultures.** Sphingolipids detected *via* FI-MS in all single- and co-cultures grouped by family. *C. perfringens* is shown separately from the *Clostridiaceae*. The intensity was normalized to the mean intensity of each metabolite across all samples. All replicates are shown in the heatmap (N = 3).

**References**

1	Kanehisa, M., Furumichi, M., Sato, Y., Kawashima, M. & Ishiguro-Watanabe, M. KEGG for taxonomy-based analysis of pathways and genomes. *Nucleic Acids Res* **51**, D587-D592, doi:10.1093/nar/gkac963 (2023).

2	Fuchs, A.-R. & Bonde, G. J. The nutritional requirements of *Clostridium perfringens*. *Microbiology* **16**, 317-329, doi:10.1099/00221287-16-2-317 (1957).

3	Goldner, S. B., Solberg, M. & Post, L. S. Development of a minimal medium for *Clostridium perfringens* by using an anaerobic chemostat. *Applied and Environmental Microbiology* **50**, 202-206, doi:10.1128/aem.50.2.202-206.1985 (1985).

4	Sebald, M. & Costilow, R. N. Minimal growth requirements for *Clostridium perfringens* and isolation of auxotrophic mutants. *Appl Microbiol* **29**, 1-6, doi:10.1128/am.29.1.1-6.1975 (1975).
